# The Irish potato famine pathogen subverts host vesicle trafficking to channel starvation-induced autophagy to the pathogen interface

**DOI:** 10.1101/2020.03.20.000117

**Authors:** Pooja Pandey, Alexandre Y Leary, Yasin Tümtas, Zachary Savage, Bayantes Dagvadorj, Emily Tan, Virendrasinh Khandare, Cian Duggan, Temur Yusunov, Mathias Madalinski, Federico Gabriel Mirkin, Sebastian Schornack, Yasin Dagdas, Sophien Kamoun, Tolga O. Bozkurt

## Abstract

Eukaryotic cells deploy autophagy to eliminate invading microbes. In turn, pathogens have evolved effector proteins to counteract antimicrobial autophagy. How and why adapted pathogens co-opt autophagy for their own benefit is poorly understood. The Irish famine pathogen Phythophthora infestans secretes the effector protein PexRD54 that selectively activates an unknown plant autophagy pathway, while antagonizing antimicrobial autophagy. Here we show that PexRD54 induces autophagosome formation by bridging small GTPase Rab8a-decorated vesicles with autophagic compartments labelled by the core autophagy protein ATG8CL. Rab8a is required for pathogen-triggered and starvation-induced but not antimicrobial autophagy, revealing that specific trafficking pathways underpin selective autophagy. We discovered that Rab8a contributes to basal immunity against P. infestans, but PexRD54 diverts a sub-population of Rab8a vesicles to lipid droplets that associate with autophagosomes. These are then diverted towards pathogen feeding structures that are accommodated within the host cells. We propose that PexRD54 mimics starvation-induced autophagy by channeling host endomembrane trafficking towards the pathogen interface possibly to acquire nutrients. This work reveals that effectors can interconnect independent host compartments to stimulate complex cellular processes that benefit the pathogen.

**Graphical abstract:** 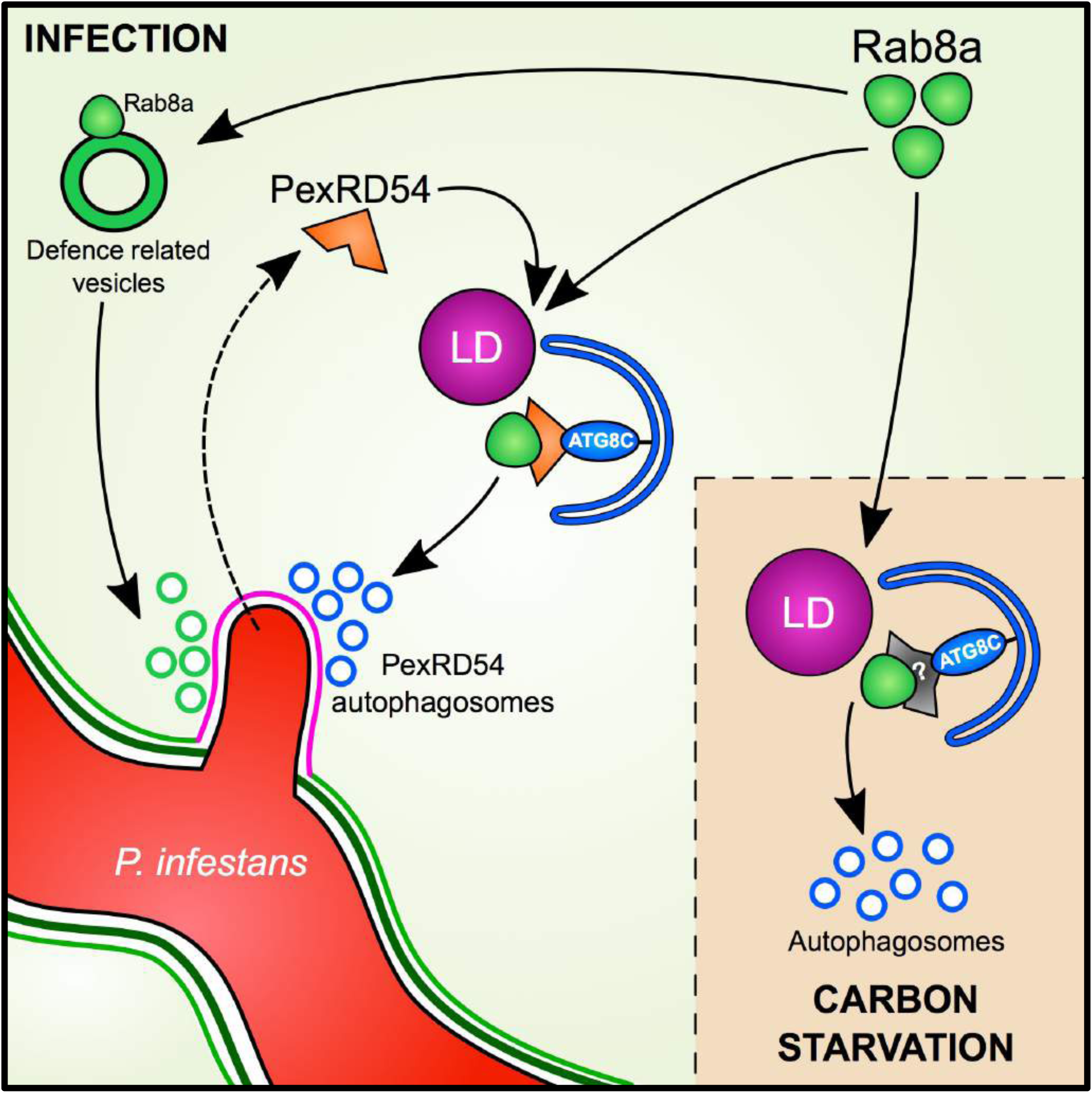

## Introduction

Autophagy is a conserved eukaryotic cellular process that mediates the lysosomal degradation and relocation of cellular cargoes within double-membraned vesicles called autophagosomes (He and Klionsky, 2009; Leidal et al., 2020). Although autophagy is historically considered to be a bulk catabolic pathway tasked with maintaining cellular homeostasis under normal or stress conditions, it is now clear that autophagy can be highly selective in cargo choice (Zaffagnini and Martens, 2016). Autophagic cargoes are typically captured during autophagosome formation, a complex process that is regulated by the concerted action of a set of conserved autophagy related proteins (ATG) as well as specialized autophagy adaptors and cargo receptors (Mizushima et al., 2011). These captured cargoes are sorted within the autophagosome during maturation of the isolation membrane (also known as the phagophore) via the specific interactions between cargo receptors and a lipidated form of ATG8, which decorates the phagophore to serve as a docking platform for cargo receptors (Ryabovol and Minibayeva, 2016; Slobodkin and Elazar, 2013).

The source of the phagophore is still under debate, but its primary source is considered to be the endoplasmic reticulum (ER) (Bernard and Klionsky, 2013; Hamasaki et al., 2013). As the cargo is being captured, the phagophore undergoes massive expansion and finally gets sealed to form a mature autophagosome. Therefore, formation of the autophagosome requires additional lipid supplies that are needed for elongation and final sealing of the phagophore. Supporting this view, the essential autophagy protein ATG2 was recently discovered to have lipid transfer activity (Maeda et al., 2019; Osawa et al., 2019; Valverde et al., 2019). To cope with cellular starvation, cells can rapidly generate hundreds of autophagosomes that conceivably require an efficient supply of lipids. Remarkably, in yeast, lipids mobilized from lipid droplets (LDs) were found to fuel autophagosome biogenesis during starvation-induced autophagy, which employs relatively larger autophagosomes. In contrast, smaller autophagosomes of the cytosol-to-vacuole transport (Cvt) pathway do not rely on LDs, suggesting that LDs are specifically recruited for starvation-induced autophagy in order to meet the increased demand of lipids required for the biogenesis of larger sized autophagosomes (Dupont et al., 2014; Shpilka et al., 2015). Although poorly characterized, there is accumulating evidence for autophagosome maturation relying on vesicle transport and membrane expansion events which are regulated by secretory pathways involving Rab GTPases (Ao et al., 2014). However, the molecular mechanisms governing autophagosome biogenesis, including the sources of membrane precursors required for autophagosome elongation and the transport routes to position these lipid supplies at autophagosome assembly sites remain poorly understood.

Discovery of an increasing number of autophagy cargo receptors uncovered a multitude of selective autophagy pathways implicated in crucial cellular functions ranging from development to immunity in both plants and animals (Zaffagnini and Martens, 2016). For instance, the plant selective autophagy (aggrephagy) cargo receptor Joka2/NBR1 mediates antiviral immunity by eliminating viral components through autophagy (Hafrén et al., 2017a, 2017b; Jung et al., 2020). Joka2/NBR1 is also required for immunity against bacteria and oomycete pathogens, however, the extent to which defense-related autophagy acts against these pathogens is unknown (Dagdas et al., 2018; Hofius et al., 2018). Consistent with the important role of autophagy in plant immunity, adapted pathogens appear to have evolved strategies to manipulate the host autophagy machinery to support their virulence (Hofius et al., 2018; Leary et al., 2017, 2019).

Pathogens typically secrete an arsenal of effector proteins to modulate host processes to support their virulence. Effectors function not only to evade and suppress host immunity but also to mediate nutrient acquisition (Win et al., 2012). Some filamentous plant pathogens, including the Irish potato famine pathogen *Phytophthora infestans*, project hyphal extensions called haustoria that grow into the host cells to facilitate effector delivery and gain access to host nutrients (Panstruga and Dodds, 2009). A haustorium is a specialized infection structure that remains enveloped by an enigmatic host-derived membrane known as the extra-haustorial membrane (EHM), whose functions and biogenesis are poorly understood (Whisson et al., 2016). Notably, we previously showed that Joka2-mediated defense-related autophagy is diverted to the EHM during *P. infestans* infection (Dagdas et al., 2018). The pathogen counteracts this by deploying PexRD54, a host-translocated RXLR class of effector with five consecutive WY motifs, that targets plant autophagy (Dagdas et al., 2016). PexRD54 carries a canonical C-terminal ATG8 interacting motif (AIM) that is typically found on autophagy cargo receptors to bind ATG8 (Maqbool et al., 2016). Among the diverse set of potato ATG8 members (Kellner et al., 2017; Zess et al., 2019), PexRD54 preferentially binds the ATG8CL isoform and outcompetes Joka2/NBR1 from ATG8CL complexes, thereby disarming defense-related autophagy at the pathogen interface. Intriguingly, PexRD54 does not fully shutdown autophagy as has been shown for animal pathogens that suppress autophagy (Choy et al., 2012; Kimmey and Stallings, 2016; Real et al., 2017; Xu et al., 2019). Instead, it stimulates formation of autophagosomes that accumulate around the pathogen interface. However, how PexRD54 stimulates autophagy and in what way the pathogen benefits from this remains unknown.

Here, we show that PexRD54 mimics carbon starvation induced autophagy by coupling the host vesicle transport regulator Rab8a to autophagosome biogenesis at the pathogen interface. While Rab8a contributes to basal immunity against *P. infestans*, PexRD54 diverts a sub-population of the Rab8a pool to trigger autophagosome formation. Thus, using an effector protein as a molecular tool, we provide insights into not only how vesicle transport processes selectively support autophagosome formation, but also how the pathogen exploits these pathways to undermine plant immunity.

## Results

### PexRD54-ATG8 binding is not sufficient for stimulation of autophagosome formation

We have previously shown that AIM mediated binding of PexRD54 to ATG8CL is essential for the activation of autophagosome formation (Dagdas et al., 2016; Maqbool et al., 2016). Therefore, we reasoned that PexRD54 could stimulate autophagosome formation by either releasing the negative regulation of ATG8CL by host autophagy suppressors or through recruiting essential host components to the autophagosome biogenesis sites. To first address whether ATG8 binding is sufficient to trigger autophagosome induction, we generated a PexRD54 truncate comprising only the C terminal AIM peptide (amino acids 350-381, hereafter AIMp), and compared its potency to stimulate autophagosome formation to the full length protein. Strikingly, instead of stimulating autophagosome formation, AIMp fused to RFP (RFP:AIMp) significantly reduced the number of ATG8CL-autophagosomes in leaf epidermal cells (Figure 1A-B). Compared to RFP:GUS control, expression of RFP:AIMp reduced the number of GFP:ATG8CL-autophagosomes by ∼6 fold, whereas cells expressing RFP:PexRD54 had a ∼4 fold increase in GFP:ATG8CL-autophagosome numbers as has been shown before (Figure 1A-B) (Dagdas et al., 2016). The AIMp interacted with ATG8CL *in planta* (Figure S1), as was previously shown through *in vitro* studies (Dagdas et al., 2016; Maqbool et al., 2016). However, this association appears to take place mainly in the cytoplasm as the suppression of autophagosome formation by AIMp is such that we hardly observe any GFP:ATG8CL autophagosomes (Figure 1B). In contrast, RFP:PexRD54 and GFP:ATG8CL produced strong overlapping fluorescence signals that peak at mobile ring-like PexRD54-ATG8CL-autophagosome clusters as described previously (Dagdas et al., 2016), in contrast to cells expressing RFP:GUS (Figure 1B, S2). Taken together these results show that binding of PexRD54 to ATG8CL, although necessary, is not sufficient to activate autophagosome biogenesis. This suggests while the full-length protein stimulates autophagy, PexRD54’s AIM peptide functions as an autophagy suppressor.

**Figure 1.**
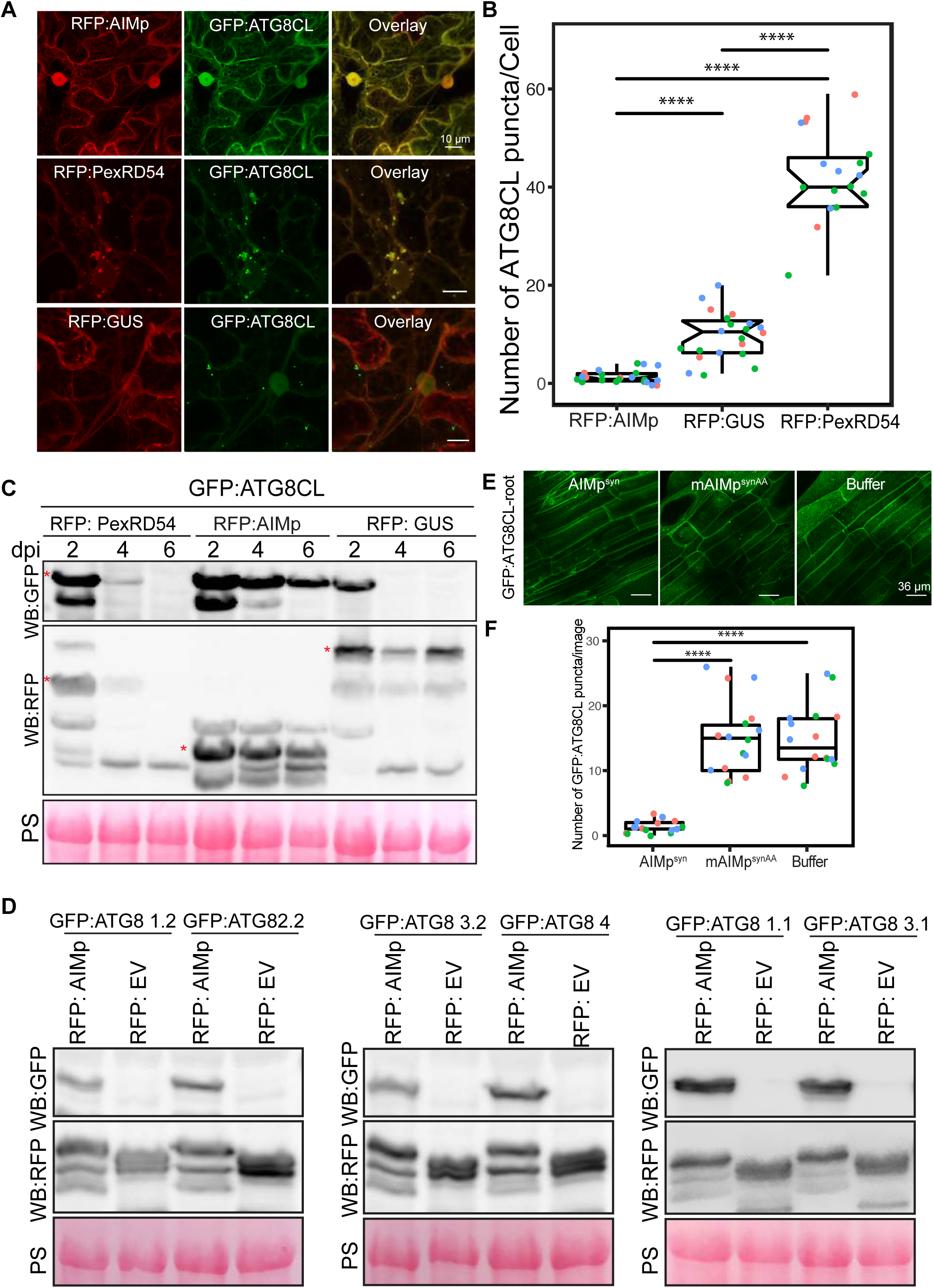
PexRD54-ATG8 binding is not sufficient for stimulation of autophagosome formation. (A) Maximum projection confocal micrographs of *Nicotiana benthamiana* leaf epidermal cells transiently expressing either RFP:AIMp (top), RFP:PexRD54 (middle), or RFP:GUS (bottom), with GFP:ATG8CL. Scale bars = 10 µm. (B) Quantification of autophagosome numbers from A shows RFP:PexRD54 expression significantly increases ATG8CL autophagosomes per cell (40, *N* = 19 images quantified) compared to RFP:GUS control (10, *N = 22* images quantified), while RFP:AIMp significantly decreases ATG8CL autophagosome numbers (2, *N* = 23 images quantified). Scattered points show individual data points, colour indicates biological repeats. (C) Western blots show Agrobacterium mediated expression of GFP:ATG8CL is stabilized by RFP:AIMp and RFP:PexRD54 beyond two days post infiltration. Red asterisks show bands of expected sizes. (D) Western blots show Agrobacterium mediated expression of various GFP:ATG8 isoforms is stabilized by RFP:AIMp. (E) Maximum projection confocal micrographs of transgenic *N. benthamiana* leaf cells stably expressing GFP-ATG8CL infiltrated with cell penetrating peptides or buffer. (F) Compared to a buffer control (15, *N* = 16 images quantified), synthesized AIM peptide fused to a cell penetrating peptide (AIMpsyn) suppresses ATG8CL autophagosomes in roots (1, *N* = 18 images quantified), while the AIM peptide mutant mAIMp^synAA^ does not (15, *N* = 17 images quantified).

### The AIM peptide of PexRD54 suppresses autophagy

To determine whether AIMp negatively regulates autophagy, we investigated its impact on autophagic flux by monitoring GFP:ATG8CL depletion over time. Consistent with the AIMp triggered decrease in ATG8CL-autophagosome numbers (Figure 1A-B), RFP:AIMp stabilized GFP:ATG8CL compared to RFP:PexRD54 or RFP:GUS control (Figure 1C, S3). Western blotting showed that in the presence of RFP:AIMp, GFP:ATG8CL was still able to produce a strong protein band even after six days post transient expression. On the other hand, GFP:ATG8CL was hardly detectable just four days in presence of RFP:PexRD54 or RFP:GUS, indicating that the AIMp hampers autophagic flux (Figure 1C). However, we did not observe any stabilization of the GFP control by RFP:AIMp, RFP:PexRD54 or RFP:GUS, indicating that reduced turnover of GFP:ATG8CL by RFP:AIMp is specific and is not due to altered Agrobacterium-mediated expression efficiency (Figure S3).

We next investigated the extent to which AIMp acts on other potato ATG8 isoforms. RFP:AIMp showed a robust stabilization effect on all six potato ATG8 isoforms (Figure 1D), indicating that it acts as a broad spectrum autophagy suppressor. To further validate these results, we measured the endogenous NBR1/Joka2 and ATG8 protein levels in *N. benthamiana* in the presence or absence of the AIMp. Consistently, ectopic expression of RFP:AIMp led to marked increase in both native NbJoka2 and NbATG8(s) levels compared to RFP:GUS expression, whereas RFP:PexRD54 mildly increased levels of only native NbJoka2 but not NbATG8s (Figure S4). These results further support the view that PexRD54’s AIMp suppresses autophagy non-selectively, whereas PexRD54 activates ATG8CL-autophagy while neutralizing Joka2-mediated autophagy as shown before (Dagdas et al., 2016).

We then explored the potency of the AIMp in autophagy suppression when applied exogenously. For this we custom synthesized PexRD54’s AIM peptide (AIMp^syn^, 10 amino acids at the C terminus) along with the AIM peptide mutant (mAIMp^syn^) fused to cell penetrating peptides and tested their activities in plant roots that stably express GFP:ATG8CL. Although both AIMp^syn^ and mAIMp^syn^ fused to 5-Carboxyfluorescein (CF-AIMp^syn^ and CF-mAIMp^syn^) were effectively taken up by the root cells (Figure S5), only the wild type AIMp^syn^ reduced the frequency of GFP:ATG8CL-puncta (by ∼10-fold) compared to mAIMp^syn^ or water control (Figure 1E-F). We also repeated these assays in leaf epidermal cells successfully, however, peptide translocation efficiency and thus autophagosome reduction AIMp^syn^ were much lower in leaves compared to root cells (Figure 1E-F, S6). These findings demonstrate that PexRD54’s AIM peptide suppresses autophagy, likely through binding to plant ATG8 isoforms with a high affinity and limiting their access to the autophagy adaptors that are essential for induction of autophagy. This hints that full-length PexRD54 carries additional features to stimulate autophagosome formation, by for instance, recruiting and/or manipulating other host components.

### PexRD54 associates with the host vesicle transport regulator Rab8a independent of ATG8CL binding

As our previous findings revealed that host autophagy function is important for *P. infestans* infection (Dagdas et al., 2016, 2018), we next set out to investigate the mechanism of autophagy activation by PexRD54. Although the underlying molecular mechanisms are largely unknown, autophagosome biogenesis relies on vesicle trafficking and fusion events in yeast and animals (Nair et al., 2011; Singh et al., 2019). We therefore reasoned that in addition to binding ATG8CL, PexRD54 could possibly hijack host vesicle transport machinery to stimulate autophagosome biogenesis. Interestingly, our previous proteomics survey identified Rab8a, a member of the small ras-related GTPases that mediate vesicle transport and fusion events, as a candidate PexRD54 interactor (Dagdas et al., 2016). We first validated PexRD54-Rab8a association through co-immunoprecipitation assays by co-expressing the potato Rab8a (herein Rab8a) with PexRD54 *in planta*. Notably, the AIM mutant of PexRD54 (PexRD54^AIM^) that cannot bind ATG8CL still interacted with Rab8a to a similar degree as PexRD54 (Figure 2A), indicating that PexRD54 associates with Rab8a independent of its ATG8CL binding activity. Consistent with this, the AIMp failed to associate with Rab8a in pull down assays, although it still strongly interacted with ATG8CL (Figure 2B). These results suggest that PexRD54’s N-terminal region preceding the C-terminal AIM mediates Rab8a association.

**Figure 2.**
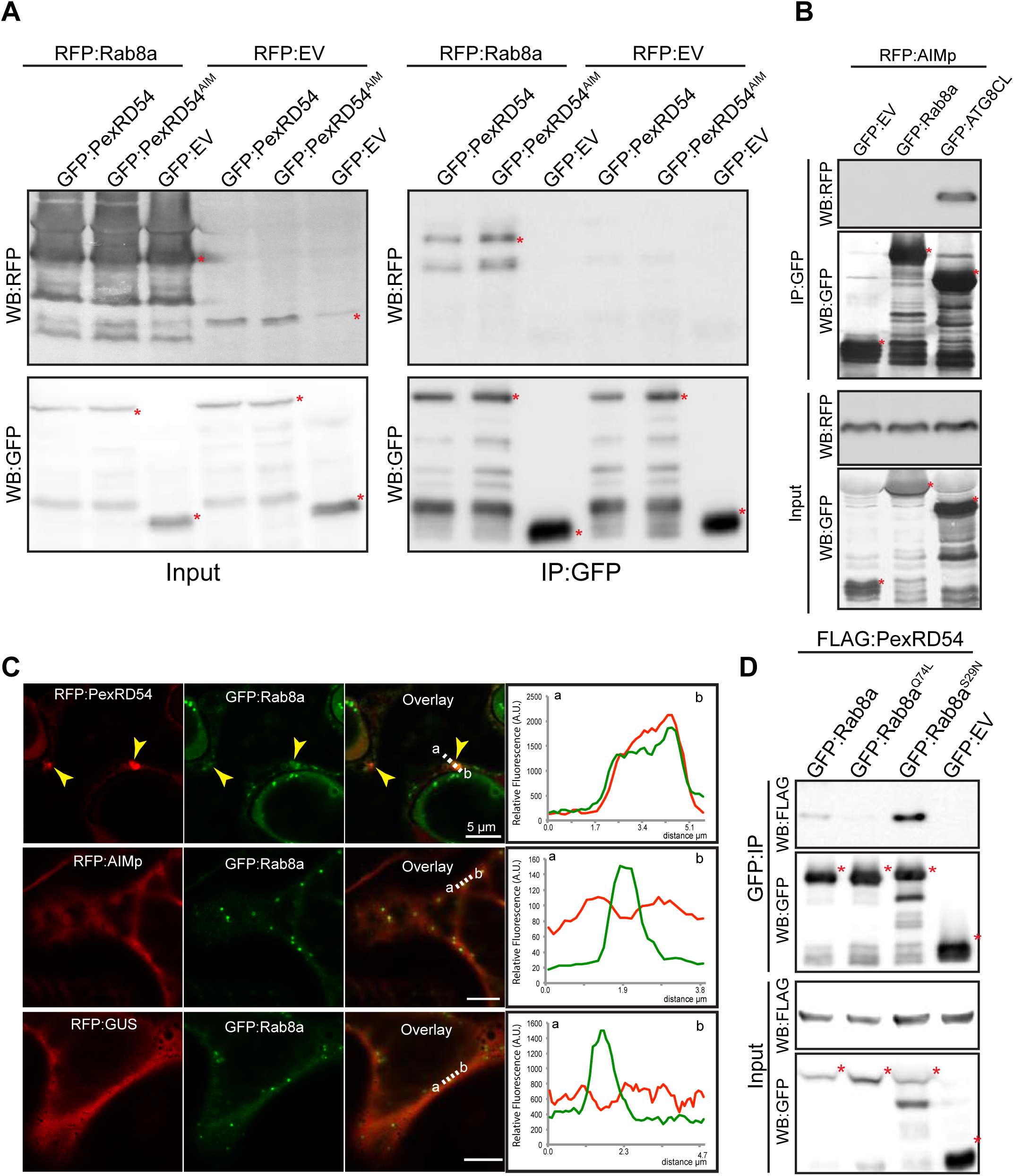
PexRD54 associates with the host vesicle transport regulator Rab8a independently of its ATG8CL binding. (A) *In planta* co-immunoprecipitation between Rab8a and PexRD54 or PexRD54^AIM^. RFP:Rab8a was transiently co-expressed with either GFP:EV, GFP:PexRD54 or GFP:PexRD54^AIM^. Red asterisks indicate expected band sizes. (B) *In planta* co-immunoprecipitation between the AIMp and AT8CL or Rab8a. RFP:AIMp was transiently co-expressed with either GFP:EV, GFP:AT8CL or GFP:Rab8a. IPs were obtained with anti-GFP antiserum. Total protein extracts were immunoblotted. Red asterisks indicate expected band sizes. (C) Maximum projection confocal micrographs of *N. benthamiana* leaf epidermal cells transiently expressing either RFP:PexRD54 (top), RFP:AIMp (middle), or RFP:GUS (bottom), with GFP:Rab8a. Yellow arrows show colocalization between constructs. Transects in overlay panel correspond to plot of relative fluorescence over the labelled distance. RFP:PexRD54 co-localises in discrete punctate structures with GFP:Rab8a while RFP:AIMp and RFP:GUS show diffuse expression. (D) *In planta* co-immunoprecipitation of PexRD54 with Rab8a, Rab8a^Q74L^, Rab8a^S29N^, or GFP. FLAG:PexRD54 was transiently co-expressed with GFP:Rab8a, GFP:Rab8a^Q74L^, GFP:Rab8a^S29N^, or GFP:EV. IPs were obtained with anti-GFP antiserum and total protein extracts were immunoblotted with GFP and FLAG antisera. Red asterisks indicate expected band sizes.

To gain insights into association of PexRD54 with Rab8a, we investigated their subcellular distribution through confocal microscopy in leaf epidermal cells. Both stably and transiently expressed Rab8a fused to GFP (GFP:Rab8a) produced fluorescent signals at both the plasma membrane and the vacuolar membrane (tonoplast) (Figure S7). In addition, GFP:Rab8a localized to mobile puncta (0.2-0.5 μm in diameter) as well as to larger ring shaped structures (Figure S7, Video S1), indicating that Rab8a could regulate multiple cellular trafficking events. To determine the subcellular compartment(s) where PexRD54 associates with Rab8a, we examined GFP:Rab8a expressed with RFP:PexRD54, RFP:AIMp or RFP:GUS using confocal microscopy. In line with the pull-down assays (Figure 2A-B), a subset of punctate structures labelled by GFP:Rab8a showed a clear overlap with RFP:PexRD54-puncta, whereas we did not detect any RFP signal peaking at GFP:Rab8a puncta in cells expressing RFP:AIMp or RFP:GUS (Figure 2C). Together, these results demonstrate that PexRD54 associates with Rab8a in an AIM independent manner and raise the possibility that Rab8a could be an important component of PexRD54 driven autophagy.

### PexRD54 shows a higher affinity towards Rab8a-S29N mutant

Because Rab GTPases function by converting between GTP and GDP bound states, we decided to generate Rab mutants that mimic the active (GTP) and inactive (GDP bound) conformations, which are helpful for characterization of the Rab GTPase functions. Although recent work challenged the applicability of these mutations (Langemeyer et al., 2014; Pfeffer et al., 2014), we reasoned that Rab8a mutants could still be useful to dissect the role of Rab8a in PexRD54 activated autophagy. To determine whether PexRD54 favors a particular form of Rab8a, we produced Rab8a point mutants that we presume to mimic the GTP (Rab8a^Q74L^) or GDP (Rab8a^S29N^) bound states (Figure S8A) and investigated their subcellular distribution (Figure S8B-D). Unlike GFP:Rab8a, which predominantly labelled the plasma membrane, GFP:Rab8a^S29N^ mutant showed an even distribution at the plasma membrane and the tonoplast (Figure S8B-C). In addition, both GFP:Rab8a^S29N^ and GFP:Rab8a marked punctate structures with varying size and shape (Figure S8B-C). In contrast, GFP:Rab8a^Q74L^ was mainly trapped in the tonoplast and showed reduced punctate distribution compared to GFP:Rab8a or GFP:Rab8a^S29N^ (Figure S8B, D), indicating that the Q74L mutant may not be representing the fully active form of Rab8a as previously reported for other Rab GTPases (Langemeyer et al., 2014; Pfeffer et al., 2014).

We next examined the extent to which Rab8a mutants colocalize with PexRD54. When co-expressed with BFP:PexRD54, both GFP:Rab8a and GFP:Rab8a^S29N^ consistently produced sharp fluorescence signals that overlap with the typical ring-like autophagosomes marked by PexRD54 (Figure S9A-B). However, GFP:Rab8a^Q74L^ showed a similar localization pattern to the GFP control, and mostly did not produce fluorescence signals that peak at BFP:PexRD54-puncta (Figure S9C-D). We quantified these observations in multiple independent experiments where GFP:Rab8a and GFP:Rab8a^S29N^ frequently (68%, N = 23) labeled BFP:PexRD54-puncta, whereas GFP:Rab8a^Q74L^ only did so much less often (25%, N = 20) (Figure S9E). As an additional control, we also checked for colocalization between Rab8a mutants and PexRD54’s AIM peptide. However, we did not observe any puncta co-labeled by RFP:AIMp and GFP:Rab8a or any of the Rab8a mutants we tested (Figure S10). These observations are consistent with the results obtained in Figure 2B and 1A-B which revealed that PexRD54’s AIM peptide fails to associate with Rab8a and suppresses autophagosome formation. Finally, we tested the affinity of PexRD54 to Rab8a and its mutants. Rab8a^S29N^ pulled-down PexRD54 more than wild type GFP:Rab8a or GFP:Rab8a^Q74L^ *in planta* (Figure 2D). This suggests that PexRD54 preferentially associates with the GDP (S29N) bound state of Rab8a.

### Rab8 family contributes to immunity against *P. infestans*

We next investigated the potential role of Rab8a in immunity against *P. infestans*. First, we tested whether silencing of Rab8a gene expression interferes with the pathogen growth. In the *N. benthamiana* genome, we identified at least four genes encoding full-length Rab8a like proteins (NbRab8a1-4). An RNA interference (RNAi) silencing construct (RNAi:NbRab8a^1-2^ hereafter) targeting the 3 prime untranslated regions (UTR) of *NbRab8a1* and *NbRab8a2*, the two closest homologs of the potato *Rab8a* in *N. benthamiana*, showed efficient silencing of the two genes but not *NbRab8a3 and NbRab8a4*, compared to the control silencing construct RNAi:GUS (Fig S11). In three independent experiments RNAi mediated silencing of two out of four Rab8a homologs did not alter *P. infestans* virulence (Figure S12). We reasoned that this could be due to redundancy among Rab8a homologues that are potentially upregulated during infection. To overcome this limitation, we generated a hairpin-silencing construct (RNAi:NbRab8a^1-4^) that targets all four Rab8a members in *N. benthamiana*. The RNAi:NbRab8a^1-4^ construct effectively silenced all four Rab8a members but not an unrelated Rab GTPase family member Rab11 (Figure S13). Simultaneous silencing of all four Rab8a members led to a consistent increase in disease symptoms caused by *P. infestans* (Figure 3A), supporting the view that the Rab8a family contributes to immunity. To determine the role of the Rab8a family in immunity, we conducted a silencing complementation assays using a codon shuffled Rab8a-1 construct fused to GFP (GFP:Rab8a-1^syn^) that can evade RNAi. As expected, the GFP:Rab8a^syn^ construct was resistant to RNAi mediated silencing, as it produced a clear protein band in western blots, unlike the wild type Rab8a that was undetectable upon co-delivery of the RNAi:NbRab8a^1-4^ silencing construct (Figure S14). The enhanced susceptibility caused by RNAi:NbRab8a^1-4^ silencing was rescued by simultaneous overexpression of GFP:Rab8a-1^syn^ but not the GFP control, providing further evidence that Rab8a is required for basal resistance against *P. infestans* (Figure 3B). Finally to gain supporting evidence of the positive role of Rab8a in immunity, we used a dominant negative form of potato Rab8a (Rab8a^N128I^) to assay for immune phenotypes. Consistent with prior experiments, overexpression of GFP:Rab8a^N128I^, but not a GFP control, enhanced plant susceptibility to *P. infestans*, supporting that the Rab8a family contributes to plant immunity (Figure 3C). Taken together, these results show that Rab8a members redundantly contribute to plant immunity.

**Figure 3.**
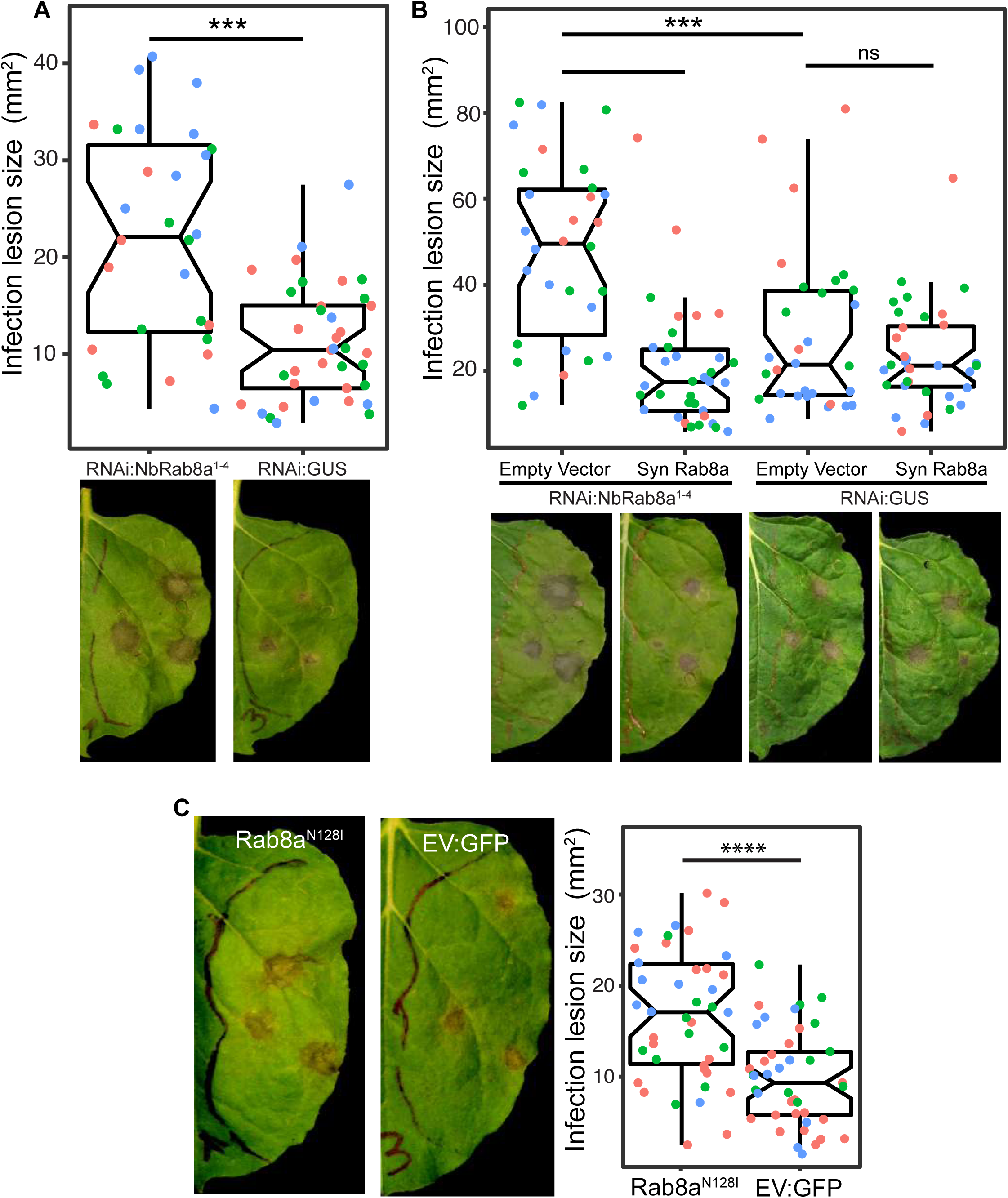
Rab8a positively contributes to immunity to *P. infestans*. (A) Silencing endogenous *Rab8a 1-4* significantly increases *P. infestans* infection lesion size (22, *N* = 28 infected leaves) compared to a silencing control (11, *N* = 37 infected leaves). *N. benthamiana* leaves expressing RNAi:NbRab8a^1-4^ or RNAi:GUS were infected with *P. infestans* and pathogen growth was determined by measuring infection lesion size 7 days post-inoculation. (B) Complementing *Rab8a 1-4* silencing with a silencing resistant GFP:Rab8a recovers resistance to *P. infestans*. When *Rab8a 1-4* is silenced, expression of SynGFP:Rab8a significantly reduces *P. infestans* infection lesion size (20, *N* = 30 infected leaves) compared to an empty vector control (61, *N* = 30 infected leaves). When a silencing control is expressed, expression of SynGFP:Rab8a does not affect *P. infestans* infection lesion size (23, *N* = 28 infected leaves) compared to an empty vector (28, *N* = 31 infected leaves). (C) Expression of Rab8a^N128I^ (36, *N* = 42 infected leaves) significantly increases P. infestans necrotic lesion size compared to an empty vector control (20, *N* = 41 infected leaves). (A-C) *N. benthamiana* leaves expressing RNAi:NbRab8a^1-4^ or RNAi:GUS, SynGFP:Rab8a or GFP:EV, GFP:Rab8a^N128I^ or GFP:EV were infected with *P. infestans* and pathogen growth was determined by measuring infection lesion size 7 days post-inoculation.

### PexRD54 recruits Rab8a to autophagosome biogenesis sites

As PexRD54 activates autophagy, we next explored the potential role of Rab8a in PexRD54 triggered autophagy. To this end, we investigated the extent to which PexRD54 associates with its two host interactors, Rab8a and ATG8CL. We first employed live-cell imaging of GFP:Rab8a and RFP:ATG8CL co-expressed in combination with either BFP:PexRD54, BFP:AIMp or BFP control. These assays revealed that BFP:PexRD54, but not free BFP or BFP:AIMp, localizes to RFP:ATG8CL puncta that are also positively labeled by GFP:Rab8a (Figure 4A-C). Furthermore, GFP:Rab8a localized to ring-shaped RFP:ATG8CL clusters triggered by BFP:PexRD54, whereas no such structures occurred in cells expressing BFP control or BFP:AIMp (Figure 4A-C). Notably, our quantitative image analysis revealed that, even in the absence of PexRD54, more than half of RFP:ATG8CL-puncta (60%, *N* = 18) are positively labelled by GFP:Rab8a (Figure 4A-D). However, BFP:PexRD54 expression significantly increased the frequency of GFP:Rab8a positive RFP:ATG8CL-puncta (85%, *N* = 18) (Figure 4A-D). Conversely, we rarely detected any fluorescent puncta that were co-labeled by GFP:Rab8a and RFP:ATG8CL in the presence of BFP:AIMp (6%, *N* = 18), which strongly suppresses autophagosome formation (Fig 4C). These results indicate that Rab8a localizes to a subset of autophagy compartments marked by ATG8CL and this is further enhanced by PexRD54. Consistently, in plants stably expressing GFP:Rab8a, we observed a similar degree of PexRD54-triggered increase in ATG8CL-Rab8a colocalization in ring-shaped ATG8CL-clusters (Fig S15), suggesting that PexRD54 might boost Rab8a recruitment to autophagic compartments. To gain biochemical evidence for PexRD54 mediated recruitment of Rab8a to ATG8CL compartments, we conducted *in planta* co-immunoprecipitation assays. Notably, the association between GFP:Rab8a and RFP:ATG8CL markedly increased in the presence of HA:PexRD54 (85%, *N = 18*) compared to HA-vector control (26%, *N = 18*), whereas GFP:Rab8a-RFP:ATG8CL interaction is slightly reduced when the two proteins are co-expressed with the HA:AIMp construct (60%, *N = 18*) (Figure S15). Altogether, these results show that PexRD54 enhances Rab8a accumulation at ATG8CL-autophagosomes.

**Figure 4.**
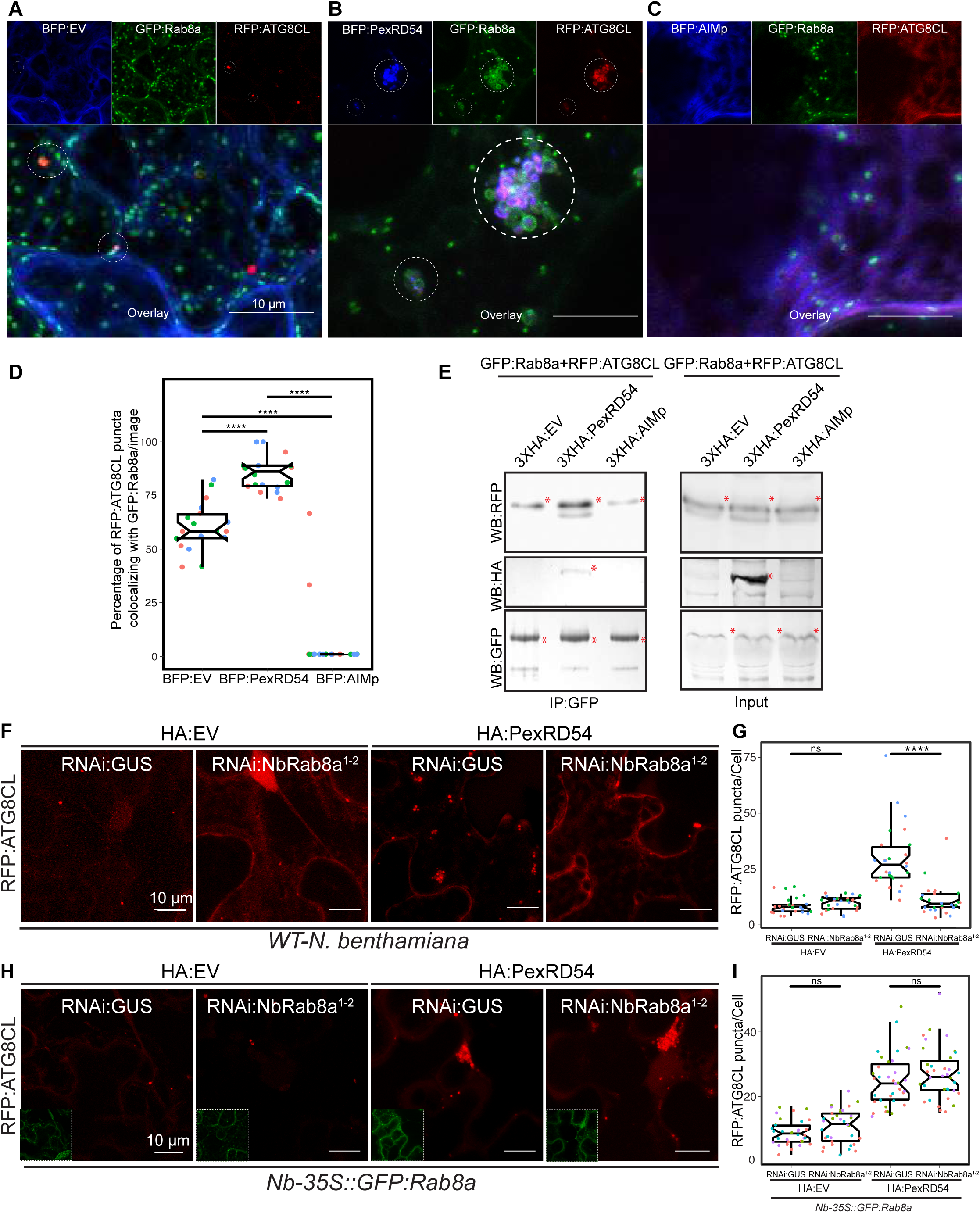
PexRD54 recruits Rab8A to ATG8CL labelled autophagosomes. (A-C) Maximum projection confocal micrographs of *N. benthamiana* leaf epidermal cells transiently expressing either BFP:EV (A), BFP:PexRD54 (B), or BFP:AIMp (C), with GFP:Rab8a and RFP:ATG8CL. Dashed white circles show variable colocalization between RFP:ATG8CL and GFP:Rab8a. (D) BFP:PexRD54 expression significantly increases punctate colocalization between RFP:ATG8CL and GFP:Rab8 (85%, *N* = 18 images quantified), while BFP:AIMp significantly reduces colocalization between RFP:ATG8CL and GFP:Rab8 (6%, *N* = 18 images quantified) compared to the BFP:EV control (60%, *N* = 18 images quantified). Scattered points show individual data points, colours indicate biological repeats. (E) *In planta* co-immunoprecipitation between Rab8A and ATG8CL, and PexRD54 or AIMp. GFP:Rab8A and RFP:ATG8CL were transiently co-expressed with either 3xHA:EV, 3xHA:PexRD54 or 3xHA:AIMp. IPs were obtained with anti-GFP antiserum. Total protein extracts were immunoblotted. Red asterisks indicate expected band sizes. (F) Confocal micrographs of *N. benthamiana* leaf epidermal cells transiently expressing RFP:ATG8CL with HA:EV or HA:PexRD54 either combined with RNAi:GUS or RNAi:NbRab8a^1-2^. (G) Silencing *N. benthamiana Rab8a1-2* significantly suppresses the autophagosome formation induced by PexRD54 (11, *N* = 26 images quantified) compared to GUS silencing control (30, *N* = 26 images quantified), but in the absence of PexRD54 silencing *Rab8a1-2* has no effect on endogenous autophagosome number (10, *N* = 27 images quantified) compared to silencing control (9, *N* = 24 images quantified). (H) Confocal micrographs of GFP:NbRab8a leaf epidermal cells transiently expressing RFP:ATG8CL with HA:EV or HA:PexRD54 either combined with RNAi:GUS or RNAi:NbRab8a^1-2^. Dashed white squares show GFP signal of complemented GFP:NbRab8a. (I) Complementing endogenous *Rab8a1-2* silencing in the silencing resistant Nb-35S::GFP:Rab8a transgenics recovered PexRD54 induced autophagosome formation (24, *N* = 35 images quantified), to similar levels to the silencing control RNAi:GUS (28, *N* = 35 images quantified).

To further ascertain the functional relationship between PexRD54 and Rab8a, we investigated the degree to which Rab8a associates with autophagy machinery. We monitored the co-localization of RFP:Rab8a with the early autophagosome biogenesis marker protein ATG9:GFP in combination with BFP:PexRD54, BFP:AIMp or BFP. Our confocal microscopy analyses revealed that ATG9:GFP puncta are frequently accompanied by RFP:Rab8a labelled vesicles. However, we detected an increased incidence of RFP:Rab8a puncta that associate with the mobile ATG9:GFP compartments in the presence of BFP:PexRD54 (68%, *N* = 44) compared to free BFP (38%, *N* = 31) or BFP:AIMp (31%, *N* = 58) (Figure S16), indicating that PexRD54 stimulates association of Rab8a with the autophagosome biogenesis machinery. Notably, only in the presence of BFP:PexRD54 but not BFP:AIMp or BFP:EV, ATG9:GFP signal overlapped with RFP:Rab8a fluorescence (Figure S16). Furthermore, time-lapse microscopy revealed that these mobile ATG9:GFP compartments co-migrate with BFP:PexRD54/RFP:Rab8a positive puncta (Video S2). These results implicate Rab8a in plant autophagy and indicate that PexRD54 promotes Rab8a recruitment to autophagosome biogenesis sites.

### Rab8a is required for PexRD54 triggered autophagy

We next investigated whether Rab8a is required for PexRD54 mediated autophagy. We measured the impact of *NbRab8a* silencing in autophagy by quantifying the RFP:ATG8CL-autophagosome numbers. For this we used the RNAi:NbRab8a^1-2^ silencing construct in order to specifically silence the *NbRab8a 1-2*. In the absence of PexRD54, silencing of *NbRab8a 1-2* did not alter the number of RFP:ATG8CL puncta (Figure S17). However, following stimulation of autophagy by transient expression of GFP:PexRD54, the number of RFP:ATG8CL-puncta/cell in RNAi:NbRab8a^1-2^ background reduced by half compared to a RNAi:GUS control (Figure S17). This suggests that simultaneous knockdown of *NbRab8a1* and *NbRab8a2* does not affect basal autophagy, but negatively impacts PexRD54 triggered autophagy. To validate these results, we set up a complementation assay in which we silenced *NbRab8a1-2* in transgenic *N. benthamiana* lines stably expressing the GFP tagged potato Rab8a, which evades RNA silencing because it lacks the 3 prime UTR targeted by the RNAi:NbRab8a^1-2^ construct. Consistent with the results obtained in Figure S17, upon delivery of RNAi:NbRab8a^1-2^ construct in wild type plants, we detected greater than two fold decrease in the number of HA:PexRD54 triggered RFP:ATG8CL-puncta compared to RNAi:GUS. On the other hand, the frequency of RFP:ATG8CL-puncta is not altered by RNAi:NbRab8a^1-2^ in cells expressing the HA vector control (Figure 4F-G). In contrast, stable transgenic plants expressing the silencing resistant potato GFP:Rab8a protein, RNAi:NbRab8a^1-2^ did not change the number of RFP:ATG8CL puncta with or without HA:PexRD54, compared to cells that express RNAi:GUS control (Figure 4H-I). These results suggest that Rab8a positively regulates PexRD54 mediated autophagy.

Finally, to gain further genetic evidence for Rab8a’s positive role in PexRD54 triggered autophagosome formation, we employed the dominant negative Rab8a mutant (N128I) (Essid et al., 2012) and measured its impact on formation of RFP:ATG8CL-autophagosomes in the presence or absence of HA:PexRD54. Consistent with the silencing assays (Figure 4F-G), GFP:Rab8a^N128I^ led to a greater than 2 fold decrease in PexRD54 triggered ATG8CL-autophagosome numbers compared to wild type GFP:Rab8a (Figure S20).

As we previously describe that PexRD54 has varying binding affinity for Rab8a and its mutants (Figure 2D), we next checked whether ectopic expression of Rab8a and its mutants (S29N and Q74L) have any effect on the formation of PexRD54-autophagosomes. Compared to GFP control, GFP:Rab8a expression led to a slight increase (∼1.5 fold) in the number of BFP:PexRD54 puncta (Figure S21), suggesting that Rab8a could positively regulate autophagosome formation. Expression of GDP bound GFP:Rab8a^S29N^ substantially boosted (∼3 times) the frequency of BFP:PexRD54 puncta compared to a GFP control, whereas GTP bound GFP:Rab8a^Q74L^ did not lead to any significant changes in the number of BFP:PexRD54 puncta compared to the GFP control (Figure S21). These results are consistent with the pulldown assays, which revealed stronger interaction between PexRD54 and Rab8a^S29N^. Together with the data presented in Figure 2D, these findings demonstrate that PexRD54 driven autophagy requires Rab8a.

### Rab8a is specifically recruited to PexRD54-autophagosomes and is dispensable for Joka2 mediated autophagy

To better characterize the autophagy stimulated by PexRD54, we decided to further investigate the interplay between Rab8a and ATG8CL. The weak interaction of ATG8CL and Rab8a in the absence of PexRD54 (Figure 4E) suggests for an indirect association potentially mediated through a host autophagy adaptor. Therefore, we explored whether increased ATG8CL and Rab8a association triggered by PexRD54 is a general hallmark of autophagy activation or is a process that is stimulated through plant selective autophagy adaptors. Because the plant autophagy cargo receptor Joka2 also binds ATG8CL and stimulates autophagosome formation (Dagdas et al., 2016), we tested if Joka2 can also stimulate Rab8a-ATG8CL association. Remarkably, unlike PexRD54, Joka2 did not interact or colocalize with Rab8a (Figure 5A, B). Moreover, our quantitative image analyses revealed that Joka2 overexpression leads to a reduction of RFP:ATG8CL puncta positively labeled by GFP:Rab8a (Figure 5C, D). This sharply contrasts with the positive impact of PexRD54 on ATG8CL-Rab8a association (Figure 5C, D), indicating that the autophagy pathway mediated by Joka2 is different from the PexRD54 triggered autophagy, and possibly does not require Rab8a function. Supporting this, we did not detect any difference in formation of Joka2 triggered autophagosomes upon *NbRab8a-1/2* silencing compared to *GUS* silencing (Fig S22).

**Figure 5.**
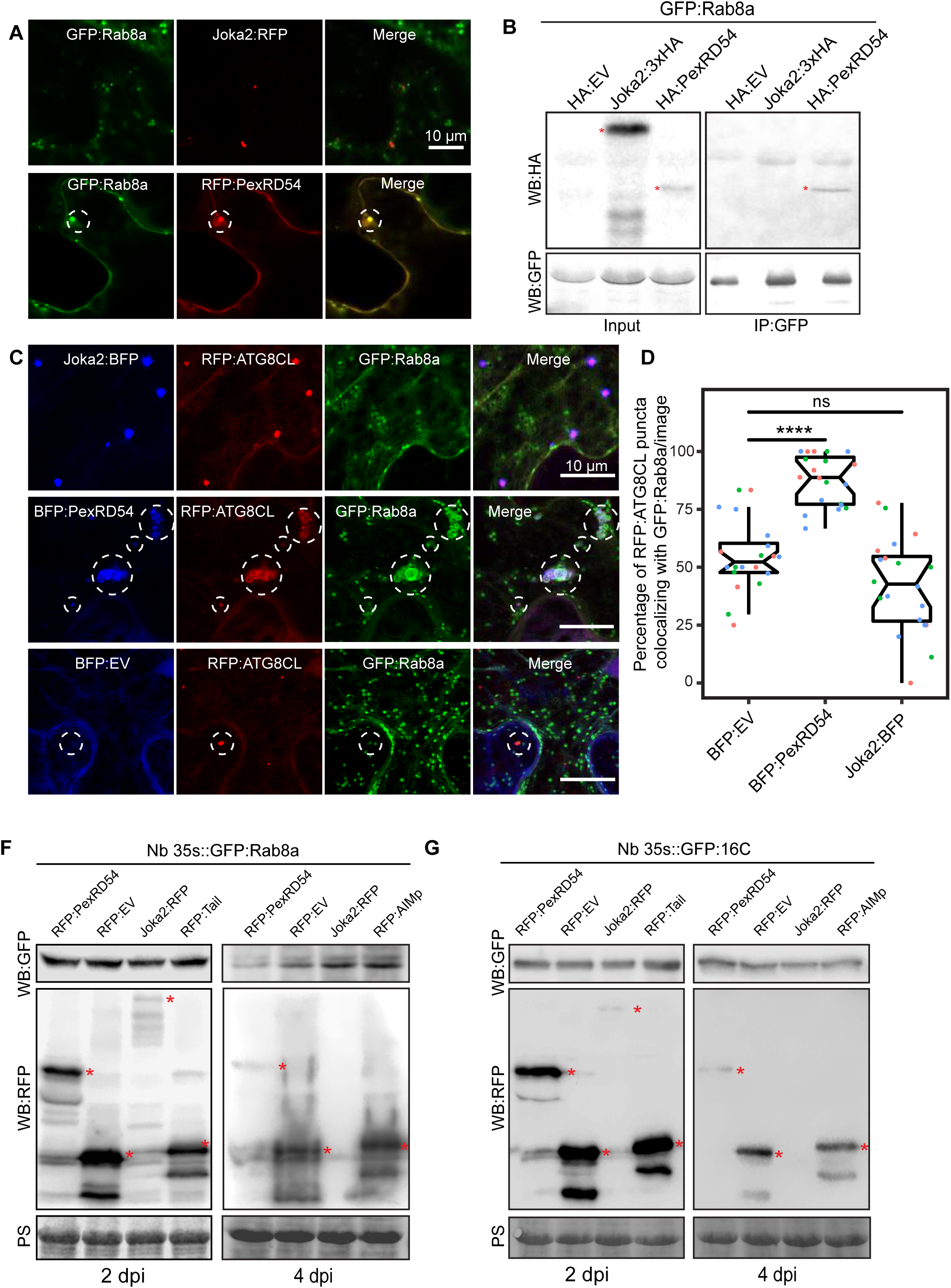
Rab8a is dispensable for Joka2-mediated autophagy. (A) Maximum projection confocal micrographs of *Nicotiana benthamiana* leaf epidermal cells transiently expressing either Joka2:RFP (top) or RFP:PexRD54 (bottom), with GFP:Rab8a. Dashed circle shows co-localized PexRD54 and Rab8a puncta. (B) *In planta* co-immunoprecipitation between GFP:Rab8a and HA:EV, Joka2:HA or HA:PexRD54. GFP:Rab8a was transiently co-expressed with HA:EV, Joka2:HA or HA:PexRD54 and immunoprecipitation was obtained with anti-GFP antiserum. Total protein extracts were immunoblotted. Red asterisks indicate expected band sizes. (C) Maximum projection confocal micrographs of *N. benthamiana* leaf epidermal cells transiently expressing either Joka2:BFP (top), BFP:PexRD54 (middle) and BFP:EV (bottom), with RFP:ATG8CL and GFP:Rab8a. (D) BFP:PexRD54 expression significantly increases colocalization of RFP:ATG8CL and GFP:Rab8a puncta (88%, *N* = 20 images quantified) compared to Joka2:BFP (42%, *N* = 20 images quantified) and BFP:EV control (55% *N* = 20 images quantified), whereas, Joka2:BFP slightly induces ATG8CL-Rab8a colocalization. Scattered points show individual data points, colour indicates biological repeat. (F) Western blotting shows that GFP:Rab8a is destabilised by expression of RFP:PexRD54 compared to RFP:EV but is stabilised by expression of Joka2:RFP or RFP:AIMp at 4dpi on *N. benthamiana* lines stably expressing the GFP tagged Rab8a. (G) Western blotting shows that GFP:EV stability is not effected by expression of RFP:PexRD54, Joka2:RFP or RFP:AIMp compared to RFP:EV at 2 or 4dpi on *N. benthamiana* lines stably expressing GFP:16C. (F-G) Proteins samples were extracted at 2 and 4 dpi and immunoblotted with GFP and RFP antisera. Red asterisks indicate expected band sizes.

Intriguingly, in stable transgenic GFP-Rab8a plants, overexpression of Joka2 led to enhanced GFP:Rab8a protein levels, in contrast to decreased GFP:Rab8a levels in presence of PexRD54 (Figure 5F). However, we did not observe any differences in GFP:EV protein levels in GFP:EV stable transgenics following ectopic expression of Joka2 or PexRD54 (Figure 5G). These findings suggest that a subset of Rab8a vesicles could be degraded by autophagy, a process that is further stimulated by PexRD54 but antagonized by Joka2. Consistent with this view, overexpression of the AIMp also led to enhanced levels of GFP:Rab8a but not GFP:EV (Figure 5F, G). Of note, the differences in GFP:Rab8a levels were only visible at later stages of ectopic expression (4 days as opposed to 2 days after transient expression) of the RFP:AIMp, RFP:PexRD54, RFP:GUS and Joka2:RFP constructs (Figure 5F-G). Collectively, these results indicate that Joka2 mediated autophagy pathway does not involve Rab8a and the weak association between ATG8CL and Rab8a observed in the absence of PexRD54 is not mediated by Joka2 but potentially through an unknown autophagy adaptor.

### PexRD54 triggers autophagy that is reminiscent of carbon starvation induced autophagy

Because autophagy can be provoked through carbon starvation, we tested whether Rab8a-ATG8CL association is altered during autophagy activation following light restriction. We detected a slight increase in the number of RFP:ATG8CL puncta upon incubation of plants for 24 hours in the dark compared to normal light conditions (Figure 6A-B). However, when GFP:PexRD54 is present, we did not measure any further enhancement of RFP:ATG8CL puncta following 24 hours in the dark, suggesting that PexRD54 mediated autophagy can override or mask starvation induced autophagy (Figure 6A-B). Furthermore, in plants stably expressing GFP:Rab8a that are exposed to 24 hours dark period, we detected an increased degree of colocalization between RFP:ATG8CL and GFP:Rab8a (Figure 6C-D, S23), in a similar fashion to enhanced ATG8CL-Rab8a association mediated by PexRD54 (Figure 4A-D). Furthermore, similar to PexRD54-mediated decrease in Rab8a levels (Figure 5F-G), we noted a reduction in GFP:Rab8a levels but not free GFP protein levels following 24 hour dark treatment of stable transgenic 35s::GFP:Rab8a plants (Figure 6E). Collectively, these data suggest that PexRD54 mimics carbon starvation induced autophagy.

**Figure 6.**
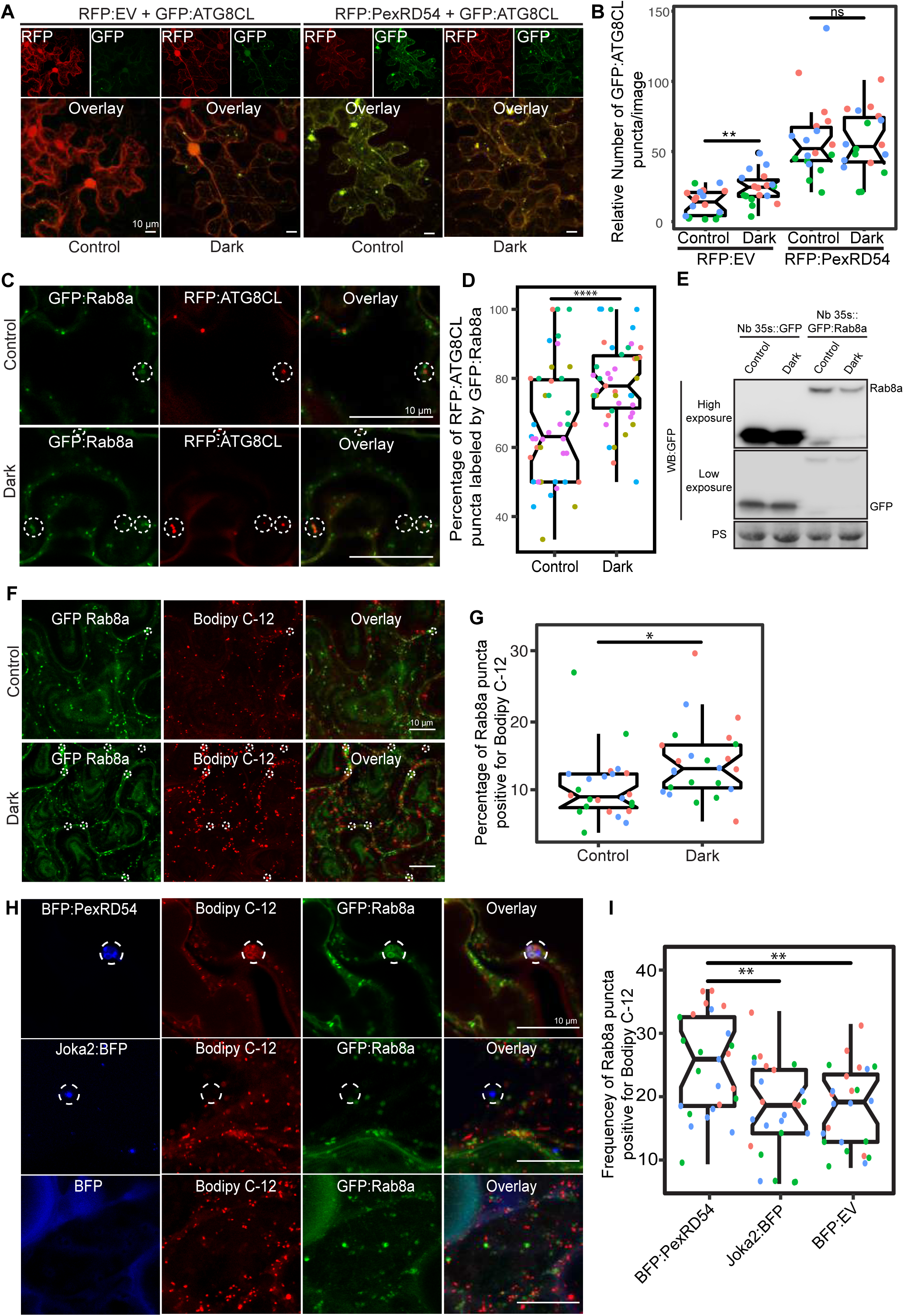
PexRD54 triggered autophagy is reminiscent of autophagy induced during carbon starvation. (A) Maximum projection confocal micrographs of *Nicotiana benthamiana* leaf epidermal cells transiently expressing either RFP:EV or RFP:PexRD54, with GFPATG8CL under normal light or 24-hour-dark conditions. (B) Scatter-boxplot shows that dark treatment (24, *N* = 18 images quantified) significantly increases RFP:ATG8CL-labelled puncta compared to control conditions (13, *N* = 18 images quantified), however when RFP:PexRD54 is present (59, *N* = 18 images quantified), dark treatment does not further enhance puncta formation (57, *N* = 18 images quantified). Images shown are maximal projections of 25 frames with 1 μm steps. Scale bars, 10 μm. (C) Maximum projection confocal micrographs of *N. benthamiana* leaf epidermal cells transiently expressing either GFP:Rab8a or RFP:ATG8CL under normal light (top) and 24-hour-dark (bottom) conditions. Dashed circle shows co-localized ATG8CL and Rab8a puncta. (D) Dark treatment significantly increases percentage of RFP:ATG8CL puncta labelled by GFP:Rab8 (79%, *N* = 44 images quantified) compared to control conditions (66%, *N* = 44 images quantified). Scattered points show individual data points, color indicates biological repeat. (E) Western blotting shows GFP:Rab8a stability is reduced by 24 hours dark unlike GFP:EV. (F) Maximum projection confocal micrographs of *N. benthamiana* leaf epidermal cells transiently expressing GFP:Rab8a and labelled by BODIPY C_12_ under normal light (top) and 24-hour-dark (bottom) conditions. Dashed circle shows co-localized Rab8a and BODIPY C_12_-positive puncta. (G) 24-hour-dark treatment increases the percentage of Rab8a puncta positive for BODIPY C_12_ (14%, *N* = 24 images quantified) compared to control conditions (10%, *N* = 24 images quantified) (H) Maximum projection confocal micrographs of *N. benthamiana* leaf epidermal cells transiently expressing either BFP:PexRD54 (top), Joka2:BFP (middle) or BFP:EV, with GFP:Rab8a and BODIPY C_12_. Dashed circle shows co-localized Rab8a, GFP-tagged PexRD54 or Joka2, and BODIPY C_12_-positive puncta. (I) Quantification of the puncta positive for both Rab8a and BODIPY C12 shows enhanced frequency of colocalization by BFP:PexRD54 (25%, *N* = 25 images quantified) but not by Joka2:BFP (18%, *N* = 24 images quantified) or BFP:EV (18%, *N* = 24 images quantified). Scattered points show individual data points, colour indicates biological repeat. Scale bars = 10 µm

Recent studies have revealed that lipid droplets (LDs) contribute to carbon starvation-induced autophagy during light restriction (Fan et al., 2019; Shpilka et al., 2015). In addition, the GDP-bound mutant form of the mammalian Rab8a is enriched at LD contact sites to regulate their fusion (Wu et al., 2014). Therefore, we decided to investigate the extent to which Rab8a and PexRD54 associate with LDs under normal or starvation conditions. We first checked co-localization between GFP:Rab8a and LDs marked by the orange-red fluorescent fatty acid (FA), BODIPY® 558/568 C12 (BODIPY-C_12_). Confocal microscopy revealed that a small fraction of GFP:Rab8a puncta are labelled by the LD marker BODIPY-C_12_ under normal light conditions, whereas the frequency of this colocalization increased by ∼1.4 fold when plants are maintained for 24 hours in the dark (Figure 6F-G). The stronger association of Rab8a and LDs upon light restriction, combined with the finding that LDs are recruited towards autophagosomes during carbon starvation (Fan et al., 2019), indicate that Rab8a-LD association could be a hallmark of starvation induced autophagy. Consistent with this view, GFP:Rab8a puncta positive for BODIPY-C_12_ are also marked by BFP:ATG8CL (Figure S24).

Strikingly, stimulation of autophagy by BFP:PexRD54, but not Joka2:BFP, led to enhanced association of BODIPY-C_12_ and GFP:Rab8a labeled puncta, supporting the hypothesis that PexRD54 mimics autophagy induced during carbon starvation (Figure 6 H-I, S25). However, we did not detect BODIPY-C_12_ fluorescence signal filling the lumen of the PexRD54 labeled compartments (Figure 6H, S27), indicating that fatty acids are likely not the autophagic cargoes of PexRD54. Rather, we detected a BODIPY-C_12_ signal at the periphery of autophagosomes marked by PexRD54, which also overlaps with GFP:Rab8a fluorescence signals (Figure 6H, S27). Moreover, these PexRD54/Rab8a-clusters are also accompanied by LDs densely labeled with only BODIPY-C_12_ as they navigate through the cytoplasm (Video S3), indicating that LDs are tightly connected to PexRD54 foci. These findings suggest that fatty acids could be one of the potential membrane sources of the autophagosomes stimulated by PexRD54 as observed during carbon starvation induced autophagy in other systems (Shpilka et al., 2015). To gain further evidence for this, we investigated the colocalization of PexRD54 and Rab8a with the LD structural membrane protein Oleosin (Fan et al., 2019; Siloto et al., 2006; Singh et al., 2009). Similar to BODIPY-C_12_ labelled puncta (Figure 6H), Oleosin labeled LDs clustered around PexRD54/Rab8a positive ring-like autophagosomes (Figure S26). Although Oleosin positive LDs were adjacent to PexRD54 autophagosomes, in contrast to BODIPY-C_12_, Oleosin-YFP did not produce fluorescent signal that overlaps with PexRD54/Rab8a ring-like autophagosomes (Figure S27), suggesting that FAs but not LD membrane proteins are transferred to PexRD54 triggered autophagosomes. Interestingly, Oleosin labeled LDs also clustered around Joka2 autophagosomes (Figure S26). However, these Oleosin clusters were not labeled with Rab8a, supporting the existence of two distinct pathways for PexRD54 and Joka2 triggered autophagy. Together these results show that PexRD54 triggers responses similar to carbon starvation-induced autophagy, including induction of autophagosomes and enhanced association of ATG8CL-autophagosomes with Rab8a and LDs.

### PexRD54 subverts Rab8a to autophagosomes at the pathogen interface

Our recent work revealed that the perihaustorial niche is a hot spot for the formation of ATG8CL autophagosomes stimulated by PexRD54 (Dagdas et al., 2016, 2018). Therefore, we next examined whether Rab8a-PexRD54 association occurs at perihaustorial ATG8CL-autophagosomes. We first checked GFP:Rab8a localization alone in the haustoriated cells. In infected leaf epidermal cells transiently or stably expressing GFP:Rab8a, we detected varying sizes of GFP:Rab8a puncta around the *P. infestans* haustoria (Figure S28). These structures included ring shaped compartments that are reminiscent of PexRD54-autophagosomes as well as smaller densely packed GFP-positive puncta and large vacuole like structures, indicating that Rab8a could regulate diverse trafficking pathways during infection (Figure S28, Video S4, S5). To verify that the perihaustorial Rab8a puncta represent the PexRD54-autophagosomes, we imaged infected plant cells which co-express GFP:Rab8a and the autophagosome marker protein RFP:ATG8CL in combination with BFP:PexRD54, BFP:AIMp, BFP or Joka2:BFP. Confocal micrographs of haustoriated plant cells showed accumulation of RFP:ATG8CL-autophagosomes around the haustoria which are co-labeled with GFP:Rab8a, and are positive for BFP:PexRD54 but not BFP control (Figure 7A). However, formation of perihaustorial puncta co-labeled by RFP:ATG8CL and GFP:Rab8a was suppressed by arresting autophagosome formation through expression of BFP:AIMp (Figure 7A). Notably, when autophagy is suppressed by BFP:AIMp or activated through BFP:PexRD54 expression, we detected a subset of perihaustorial GFP:Rab8a puncta that are not labeled by BFP:PexRD54 or RFP:ATG8CL, further supporting the view that Rab8a can mediate disparate haustorial trafficking routes (Figure 7A). We also observed that Rab8a and PexRD54 co-localize with both Bodipy C-12 and oleosin in clusters of vesicles around haustoria, linking Rab8a’s emerging role in lipid trafficking with the pathogen effector and haustorial interface (Figure S29). On the other hand, in line with our findings that the Joka2 pathway does not employ Rab8a (Figure 5), perihaustorial Joka2:BFP/RFP:ATG8CL puncta and GFP:Rab8a puncta were exclusive to each other (Figure 7A). Finally, consistent with our pull down assays (Figure 4E), we did not detect any sharp GFP:Rab8a^Q74L^ signal at the perihaustorial BFP:PexRD54 puncta (Figure S30). In contrast, GFP:Rab8a, and particularly GFP:Rab8a^S29N^, produced strong fluorescence signals peaking at perihaustorial BFP:PexRD54 puncta (Figure S30), indicating that both wild type Rab8a and Rab8a^S29N^ are enriched at the perihaustorial PexRD54 autophagosomes. These results indicate the PexRD54 stimulates diversion of Rab8a positive LDs to perihaustorial autophagosomes.

**Figure 7.**
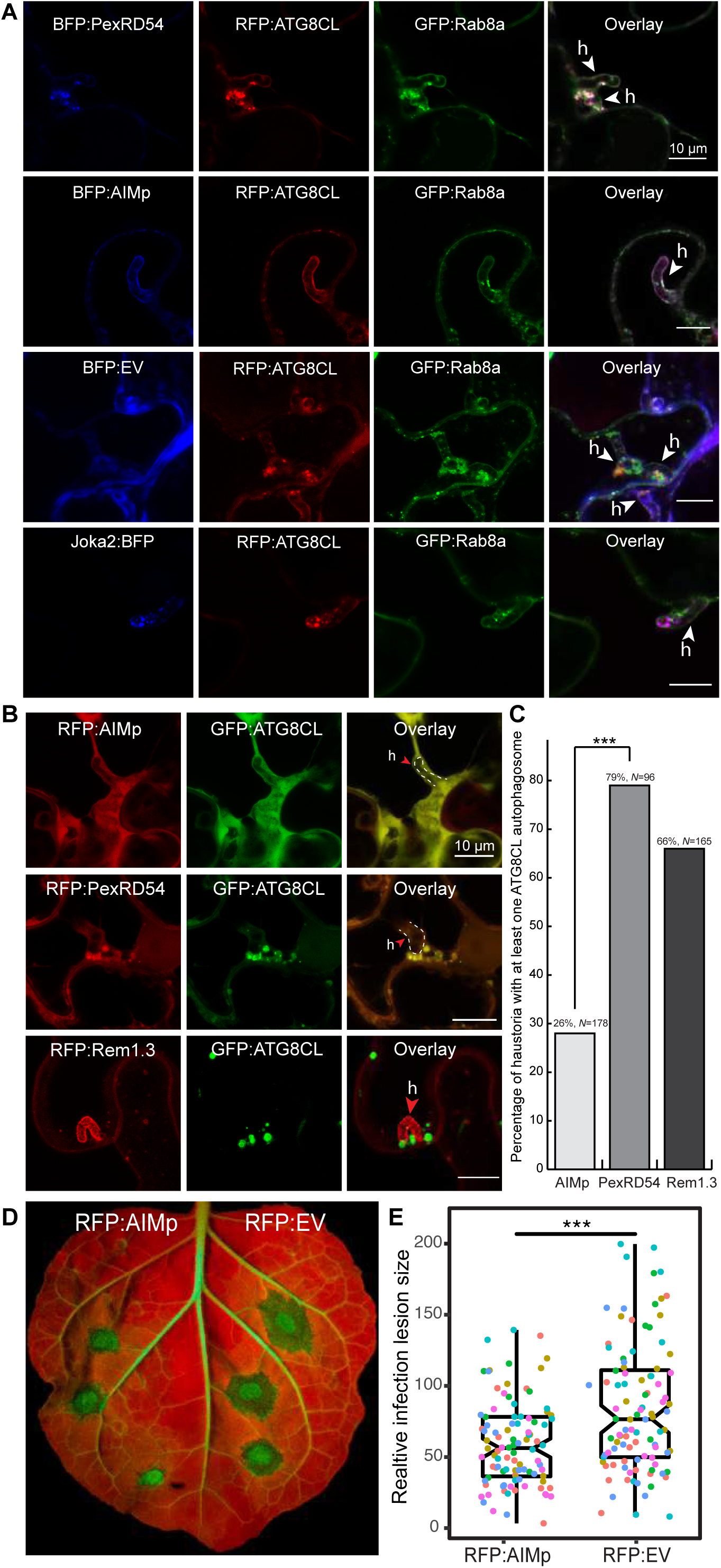
AIM peptide mediated arrest of ATG8 autophagy negatively affect *P. infestans* infection. (A) Maximum projection confocal micrographs of *P. infestans*-infected *N. benthamiana* leaf epidermal cells transiently expressing either BFP:PexRD54, BFP:EV, BFP:AIMp or Joka2:BFP, with both RFP:ATG8CL and GFP:Rab8a. BFP:PexRD54 co-localizes with RFP:ATG8CL and GFP:Rab8a at perihaustorial region, whereas, Joka2:BFP-labelled ATG8CL puncta are exclusive to GFP:Rab8a puncta. Haustoria are pointed by white arrowheads. (B) Maximum projection confocal micrographs of *P. infestans*-infected *N. benthamiana* leaf epidermal cells transiently expressing either RFP:AIMp (top), RFP:PexRD54 (middle), or PM-haustorial marker RFP:Rem1.3 (bottom), with GFP:ATG8CL. Haustoria are labelled with white dashed lines and indicated by red arrow. (C) Co-expressing RFP:PexRD54 with GFP:ATG8CL increases the percentage of haustoria associated with ATG8CL-labelled puncta (79%, *N* = 96 haustoria), compared to haustorial marker control RFP:Rem1.3 (66%, *N* = 165 haustoria) and autophagy inhibitor RFP:AIMp (26%, *N* = 178 haustoria). (D) AIMp reduces disease symptoms of *P. infestans* (65, *N* = 84 infected leaves) compared to empty vector control (87, *N* = 85 infected leaves). *N. benthamiana* leaves expressing RFP:AIMp and RFP:EV were infected with *P. infestans* and pathogen growth was determined by measuring infection lesion size 7 days post-inoculation. (E) Box plot shows relative infection lesion size of 84 and 85 infection sites, respectively from 5 biological replicates. Scattered points indicate individual data points and different colours represent various biological repeats.

### AIM peptide mediated autophagy arrest leads to reduced pathogen growth

Induction of autophagosomes by PexRD54 that are targeted to the perihaustorial interface prompted the hypothesis that *P. infestans* could benefit by co-opting host autophagy to support its own growth. Thus, we explored the impact of autophagy activation by PexRD54 on *P. infestans* host colonization. We decided to transiently interfere with pathogen induced autophagy by expressing the AIMp. Our quantitative image analysis revealed that compared to RFP:PexRD54 expression, transient expression of RFP:AIMp led to ∼3 fold decrease in the number of haustoria that are associated with autophagosomes marked by GFP:ATG8CL (Figure 7B-C). We then measured how autophagy suppression by the AIMp affects *P. infestans* infection. In multiple independent experiments (6 biological replicates), *N. benthamiana* leaf patches expressing RFP:AIMp showed a consistent reduction of quantitative disease symptoms compared to an RFP vector control (Figure 7D-E). This indicated that AIMp mediated arrest of host autophagy negatively impacts *P. infestans* virulence, supporting the hypothesis that PexRD54 triggered autophagy is beneficial to the pathogen. Collectively, these results suggest that *P. infestans* relies on host autophagy function to support its virulence. This could explain why the pathogen deploys full length PexRD54 that can activate specific host autophagy pathways while subverting defense-related autophagy, instead of just the AIM peptide.

## Discussion

Dissecting the specialized functions and mechanisms of autophagy in host-microbe interactions has been challenging. This is mainly due to prolonged stress accumulation in autophagy mutants and non-autophagy related roles of the targeted genes (Munch et al., 2014). Pathogen effectors that target specific components of the host autophagy machinery have emerged as alternative tools to unravel the underlying mechanisms of defense-related autophagy (Dagdas et al., 2016; Hafrén et al., 2017b; Hofius et al., 2018). Here, by studying the *P. infestans* effector protein PexRD54, we shed light on the poorly understood mechanism of pathogen-induced autophagy in plants. We unveil a distinct pathogen virulence mechanism in which the effector protein couples host vesicle transport machinery to autophagosome biogenesis. Through its modular domain architecture, PexRD54 employs a diverse set of host proteins such as Rab8a and ATG8CL to stimulate autophagy that is reminiscent of starvation-induced autophagy. We propose a model in which PexRD54 mimics host carbon starvation conditions to divert a sub fraction of Rab8a from defense-related vesicles to stimulate autophagosome formation. Both PexRD54 and carbon starvation drive association of Rab8a and LDs that are positioned at autophagy compartments marked by ATG8CL. Thereby, the pathogen benefits from this process through subverting the immune function of Rab8a and gaining access to plant resources carried in autophagosomes diverted to haustoria.

### AIM peptide suppressor of autophagy

Effectors have been excellent tools to dissect complex biological processes such as autophagy as effectors often have more specificity towards their targets compared to their chemical alternatives. Here, we expanded our knowledge of PexRD54 triggered autophagy and discovered an effector derived peptide that can specifically block autophagy. Chemical inhibitors of autophagy are often used to measure autophagy flux and to overcome the limitations of standard genetic approaches. However, these inhibitors are mostly inefficient and lack the required specificity. We discovered that PexRD54’s AIM peptide is a strong autophagy suppressor effective against all potato ATG8 isoforms. Conceivably, the AIM peptide competitively inhibits autophagosome biogenesis by occupying the W and L pockets on ATG8 that mediate docking of host autophagy adaptors and regulators required for autophagy activation (Noda et al., 2008). The AIM peptide is a genetically encodable tool, which can enable spatio-temporal arrest of autophagy when expressed under inducible or tissue specific promoters. Thus, this should be of great interest for autophagy studies in plants and other systems, which can overcome the limitations of chemical autophagy inhibitors and autophagy mutants. Furthermore, we developed AIM peptide derivatives with cell penetrating features, which allow studying the tissue specific functions of autophagy. The cell penetrating AIM peptide can be employed to study autophagy in plants and other eukaryotic systems, which are not amenable to genetic manipulation.

### How does PexRD54 activate autophagy?

Autophagosome biogenesis is a complex multi-step process. But how does PexRD54 activate such an intricate process? We uncovered that PexRD54 either directly or indirectly recruits Rab8a to autophagosome biogenesis sites (Figure 2-4). Similar channeling of Rab8a to ATG8CL-autophagosomes occurs during carbon starvation (Figure 4), suggesting that PexRD54 could mimic a host autophagy adaptor that regulates autophagy under nutrient deprivation. Interestingly, the mammalian autophagy cargo receptor Optineurin also interacts with both LC3 (mammalian ATG8 isoform) and Rab8a in mice (Bansal et al., 2018; Vaibhava et al., 2012). Although the functional implications of these interactions are unclear, a proposed model suggests that Optineurin mediates pre-autophagosomal membrane elongation through anchoring Rab8a to autophagosome assembly sites. Our data shows that PexRD54-ATG8CL binding is not sufficient for stimulation of autophagy (Figure 1). Therefore, it is likely that PexRD54 associates with various host components including Rab8a to drive autophagosome formation.

### The role of Rab8a in autophagy

The membrane elongation step of autophagosome formation relies on direct transport of lipids from various donor compartments. For instance, LDs provide a membrane source for autophagosome biogenesis specifically during starvation-induced autophagy (Shpilka et al., 2015). More recently, a conserved acyl-CoA synthetase (ACS) from yeast was shown to be mobilized on nucleated phagophores where it locally mediates transport of fatty acids (FAs) required for phagophore elongation (Schütter et al., 2020). But how these lipid sources such as LDs and ACS are mobilized to autophagosome biogenesis sites at first? Intriguingly, the GDP bound form of the mammalian Rab8a was found to mediate LDs fusion events (Wu et al., 2014). Furthermore, PexRD54 recruits Rab8a to autophagosome biogenesis sites and enhances Rab8a-LD localization around the ATG8CL-foci (Figure 4, 6). This suggests that PexRD54 could employ Rab8a to position LDs at ATG8CL nucleation sites to facilitate lipid transfer for autophagosome biogenesis. Interestingly, PexRD54 triggered association of Rab8a and LDs is also enhanced by carbon starvation but not by Joka2 (Figure 5, 6). We propose that upon carbon deprivation, the plant uses LDs as an alternative membrane source to accommodate for an increase in autophagosome biogenesis, and PexRD54 exploits this process to stimulate autophagy. However, Rab8a does not appear to be engaged in all autophagy routes, as it is dispensable for Joka2 mediated autophagy (Figure 6). This is consistent with the finding that Arabidopsis NBR1 (Joka2) mutants are sensitive to a variety of abiotic stress conditions but not to carbon starvation (Zhou et al., 2013). Our results are also in line with the recent findings in yeast, where LDs specifically contribute to starvation induced autophagy (Shpilka et al., 2015). Altogether, combined with previous findings, our results indicate that distinct cellular transport pathways feed autophagosome formation during diverse selective autophagy pathways in plants.

### The role of Rab8a in immunity

Our data revealed that the Rab8a family contributes to basal resistance against *P. infestans* (Figure 3). Rab8a belongs to the Rab8 family of small GTPases that are implicated in polarized secretion events in eukaryotes (Pfeffer, 2017). Another Rab8 member known as RabE1D was found to contribute to bacterial resistance and regulate membrane trafficking in the model plant Arabidopsis (Speth et al., 2009; Zheng et al., 2005). However, the extent to which Rab8 family members function in immunity remains unknown. Our findings revealed that Rab8a is most likely involved in a diverse range of cellular transport pathways including autophagy. Consistent with the evolutionarily conserved role of Rab8 family members in polarized secretion, we detected Rab8a labeling of the EHM as well as Rab8a positive vesicles around the haustorium. Interestingly, some perihaustorial Rab8a puncta were not labeled by ATG8CL or PexRD54, further supporting the view that Rab8a is engaged in diverse trafficking pathways. However, further research is required to determine the cargoes and the potential defense-related functions of the Rab8a labeled vesicles.

### Why does *P. infestans* activate autophagy?

Suppression of host autophagy by the AIM peptide leads to reduced pathogen virulence (Figure 7), indicating that suppressing host autophagy is not favorable to *P. infestans*. This could explain why the pathogen deploys PexRD54, which can neutralize defense-related autophagy mediated by Joka2, while enabling other autophagy pathways that are somewhat beneficial to the pathogen. Intriguingly, the autophagy pathway primed by PexRD54 resembles autophagy induced under carbon starvation. This hints at the possibility that PexRD54 could facilitate nutrient uptake from the host cells by mimicking starvation to stimulate the host autophagy machinery. This is further supported by our earlier finding that PexRD54-autophagosomes are diverted to the haustorial interface (Dagdas et al., 2018). PexRD54 likely remodels the autophagic cargoes engulfed in ATG8CL-autophagosomes targeted to pathogen interface, which are subsequently assimilated by the parasite. Alternatively, but not mutually exclusively, PexRD54 could also help neutralize defense-related host components by engulfing them in secure membrane-bound autophagy compartments. For instance, PexRD54 could promote diversion of Rab8a to the autophagy pathway to pacify the non-autophagy related immune functions of Rab8a.

In sum, our findings demonstrate that to support its virulence, *P. infestans* manipulates plant cellular degradative and transport systems by deploying an effector protein that imitates carbon starvation conditions. It also demonstrates effectors can act as adaptors to bridge multiple host components to modulate complex cellular processes for the benefit the pathogen. Further research is needed (i) to determine the cargo of these autophagosomes, (ii) whether they are secreted to the pathogen interface and subsequently absorbed by the pathogen, and (iii) the molecular players involved in pathogen subverted autophagy.

## Acknowledgments

We would like to thank all members of the Bozkurt Lab for their helpful discussion and suggestions. We also are grateful to the Imperial College Facility for Imaging by Light Microscopy (FILM) at Imperial College London for their imaging expertise and technical assistance. FILM is part-supported by funding from the Wellcome Trust (104931/Z/14/Z) and BBSRC (BB/L015129/1). This project was funded by the Biotechnology Biological Sciences Research Council (BBSRC) (BB/M002462/1 and BB/M011224/1), the Gatsby Charitable Foundation (GAT3395/GLD), the European Research Council, the Agencia Nacional de Promoción Científica y Tecnológica (ANPCyT, Argentina), the Royal Society (UF160413) and the Austrian Academy of Sciences.

## Methods

### Plant Material and Growth Conditions

*N. benthamiana* WT and transgenic plants (35S::GFP:Rab8a and 35S::GFP:ATG8CL) were grown and maintained in a greenhouse with high light intensity (16 hours light/8 hours dark photoperiod) at 22-24°C. To apply carbon stress, plants were kept under a dark period of 24 hours before images were acquired. Images were acquired 3 days after infiltration (dpi). 35S::GFP:Rab8a and 35S::GFP:ATG8CL lines were produced as described elsewhere (1985) with the pK7WGF2::Rab8a and pK7WGF2::ATG8CL constructs, respectively.

### Pathogenicity Assays

*P. infestans* 88069 strain was used in this study. Cultures were grown and maintained by routine passing on rye sucrose agar medium at 18°C in the dark (van West et al, 1998). Zoospores were collected from 10-14 days old culture by flooding with cold water and incubation at 4°C for 90-120 minutes. Infection of agroinfiltrated leaves was carried out by addition of 10-mL droplets of zoospore solution at 50,000 spores/ml on detached *N. benthamiana* leaves (Chaparro-Garcia et al., 2011). Infection for microscopic experiments carried out on attached leaves. Inoculated detached leaves or plants were kept in humid conditions. Day light/UV images were taken at 7 days post infection and lesion areas were measured in ImageJ.

### Virus Induced Gene Silencing (VIGS)

Virus induced gene silencing of Joka2 was performed in *N. benthamiana* as described previously (Dagdas et al., 2016). Suspensions of *Agrobacterium tumefaciens* was prepared carrying TRV1 and the TRV2:JOKA2 and mixed to a final OD600 of 0.4 and 0.2 respectively, in agroinfiltration buffer supplemented with 100 μM acetosyringone (Sigma). The suspension of agrobacterium was left in the dark for 2 h prior to agroinfiltration to stimulate virulence. TRV2:EV was used as a control. 12 days old N. benthamiana seedlings were infiltrated in cotyledons and any true leaves that had appeared. TRV2 containing the *N. benthamiana* sulphur (Su) gene fragment (TRV2-NbSU) was used as a positive control to indicate viral spread. Plants were used 4 weeks later for the *P. infestans* infection.

### Molecular Cloning and Plasmid Constructs

Various constructs used in this study were published previously. GFP:ATG8CL, GFP:PexRD54, GFP:PexRD54aim, RFP:Rem1.3 constructs were previously described in (Bozkurt et al., 2015). JOKA2:BFP, BFP:EV, BFP:PexD54, BFP:ATG8CL, ATG9:GFP, 3xHA:EV, JOKA2:3xHA constructs were described in (Dagdas et al., 2016). GFP:ATG8 1.1, GFP:ATG81.2, GFP:ATG8 2.2, GFP:ATG8 3.1, GFP:ATG8 3.2, GFP:ATG8 4 were described in (Zess et al., 2019).

RFP:PexRD54, RFP:AIMp, RFP:ATG8CL, RFP:Rab8a, Joka2:RFP, BFP:AIMp, 3xHA:PexRD54, 3xHA:PexRD54aim and 3xHA:Rd54AIMp constructs were generated by Gibson assembly of each gene PCR fragment into EcoRV digested RFP/BFP/HA vectors (N-terminal fusion for PexRD54, PexRD54aim, PexRD54AIMp and ATG8CL, C-terminal fusion for Joka2). For YFP:Oleosin, the eYFP fluorophore was split into N-terminal (residue M1-A155) and C-terminal half (residue D156 - K239). The N-terminal split YFP half was used via a linker peptide RPACKIPNDLKQKVMNH and the C-terminal split YFP half via a linker peptide HNMVKQKLDNPIKCAPR. EcoRV restriction site was added at the end of each linker to allow linearization of the vector and provide an insertion site for subsequent cloning. The DNA fragment encoding Oleosin, together with the linker peptides and restriction sites were amplified from *N. benthamiana* cDNA using primers NYFP-Oleosin F and Oleosin-CYFP R then assembled into pK7WGF2 vector backbone by Gibson assembly. GFP:Rab8a and RFP:Rab8a constructs were generated by PCR amplification from *Solanum tuberosum* cDNA using primers GW_StRab8a-1_F GW_StRab8a-1_R followed by Gateway cloning into the entry vector pENTR/D/TOPO (Invitrogen) then into the pK7WGF2 (GFP) and pH7WGR2 (RFP) vectors, respectively. RFP:GUS was created from the pENTR™-GUS control plasmid provided in the GATEWAY cloning kit and inserted into pH7WGR2 (RFP) via LR reaction. Single residue mutations of Rab8aS29N, Rab8aQ74L and Rab8aN128I were obtained by inverse polymerase chain reaction (PCR) amplification of the StRab8a entry clone with the primer pairs (phosphorylated at 5 prime ends) carrying desired mutations; (i) Rab8aS29N_F and Rab8aS29N_R; (ii) Rab8aQ74L_F and Rab8aQ74L_R; (iii) Rab8aN128I_F and Rab8aN128I_R. Templates were then eliminated by one-hour Dpn-I (New England Biolabs) restriction digestion at 37°C and the PCR products of mutants were ligated using standard protocols to obtain circular Gateway entry clones carrying desired mutations. Next, the entry clones of Rab8a mutants were recombined into destination vectors pK7WGF2 or pB7RWG2 by Gateway LR reaction. All remaining constructs were amplified from existing constructs previously described (Bozkurt et al., 2015; Dagdas et al., 2016, 2018), using primer pairs GA_RD54_F with GA_RD54_R for PexRD54, GA_RD54_F with GA_LIR2_R for PexRD54aim, GA_AIMp_F with GA_LIR2_R for PexRD54AIMp, GA_ATG8C_F with GA_ATG8C_R for ATG8CL and GA_NbJoka2_1_Fr with GA_NbJoka2_1_Rv. Silencing constructs for Rab8a were amplified using primer combinations NbRab8A_silF1 and NbRab8A_silR1, Rab8a1-4^RNAi^_F1, Rab8a1-4^RNAi^_F2, Rab8a1-4^RNAi^_F3, Rab8a1-4^RNAi^_R1, Rab8a1-4^RNAi^_R2 and Rab8a1-4^RNAi^_R3, and cloned into the pRNAiGG vector as described in Yan et al., 2012. Silencing of Rab8a was verified using RT-PCR. All primers used in this study are listed in Table 1.

**Table 1.**
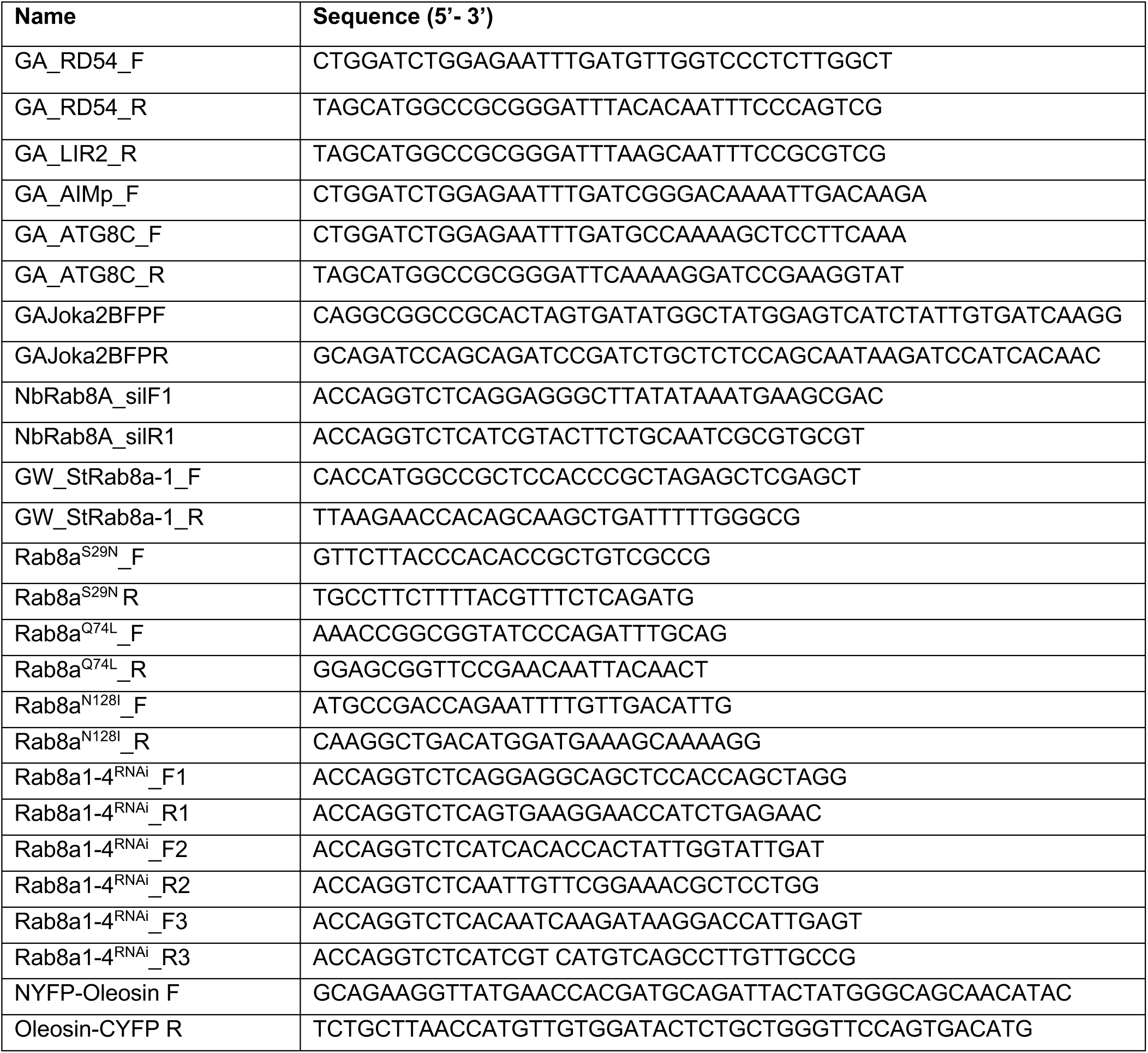
Primers used in this study.

### Co-Immunoprecipitation experiments and Immunoblot Analysis

Proteins were transiently expressed by agroinfiltration in *N. benthamiana* leaves and harvested 2 days post agroinfiltration. Protein extraction, purification and western blot analysis steps were performed as described previously (Bozkurt et al., 2011; Dagdas et al., 2016). Polyclonal anti-GFP and anti-BFP (tRFP) antibody produced in rabbit (Chromotek, UK), monoclonal antibody anti-RFP produced in mouse and monoclonal antibodies anti-GFP, anti-HA and anti-FLAG antibodies produced in rat (Chromotek, UK) were used as primary antibodies. For secondary antibodies anti-mouse antibody (Sigma-Aldrich, UK), anti-rabbit (Sigma-Aldrich, UK) and anti-rat (Sigma-Aldrich, UK) antibodies were used.

### RNA isolation, cDNA synthesis and RT-PCR

For total RNA extraction 100mg of leaf tissue excised from 3/4 days after silencing and frozen in liquid nitrogen. RNA was then extracted from the plant using GeneJET Plant RNA Purification Kit (Thermo Scientific). RNA concentration was measured using NanoDrop™ Lite Spectrophotometer (Thermo Scientific). 1 μg of RNA was used for cDNA synthesis using SuperScript IV Reverse Transcriptase (Invitrogen).

### Confocal Microscopy and Image Processing

Microscopy analyses were carried out on live leaf tissue 3-4 days post agroinfiltration. To minimize the damage of live tissue, leaf discs of *N. benthamiana* were cut using a cork borer and mounted onto Carolina observation gel (Carolina Biological Supply Company). For BODIPY-dodecanoic acid (BODIPY-C12, Invitrogen) staining, 10 μM was infiltrated into the leaf tissue 5 hours prior to observation. For PexRD54 AIM peptide experiments in leaf tissue, a solution of 10 μM of peptide in agroinfiltration buffer or buffer alone was infiltrated in leaves 3 hours prior to observation. For imaging in roots, seedlings were collected at 3 weeks old and the roots placed in 2 mL tubes containing 5 μM peptide solution in agroinfiltration buffer or buffer alone for 3 hours prior to observation.

Confocal florescence microscopy was performed using Leica SP5 and SP8 resonant inverted confocal microscope (Leica Microsystems) using 63x and 40x water immersion objective, respectively. In order to excite fluorescent tagged proteins, Diode laser excitation was set to 405 nm, Argon laser to 488 nm and the Helium-Neon laser to 561 nm and their fluorescent emissions detected at 450-480, 495-550 and 570-620 nm to visualize BFP, GFP and RFP fluorescence, respectively. Sequential scanning between lines was done to avoid spectral mixing from different fluorophores and images acquired using multichannel. Maximum intensity projections of Z-stack images were processed using ImageJ (2.0) to enhance image clarity.

### Data analysis and statistics

Images for quantification of autophagosome numbers were obtained from Z stacks consisting of 1.3 mm depth field multi-layered images with similar settings for all samples. To detect and quantify punctate structures in one channel (green channel or red channel or blue channel) and to validate colocalization an overlay of two or three channel, where applicable, was acquired (green channel and/or red channel and/or blue channel). Z stacks were separated into individual images with the ImageJ (2.0) program and analysed. To avoid possible bias of manual count, the quantification of puncta and colocalization of puncta was done using MATLAB (2017b) and ImageJ (2.0). Boxplots were generated with mean of punctate numbers generated from stacks obtained in 3-6 independent biological experiments. Statistical differences were analysed by Welch Two Sample t-test in R. Measurements were significant when p<0.05 (*) and highly significant when p<0.001(***).

### Automated puncta counting algorithm through image processing

The image processing algorithm calculates the gradient of the image to identify the boundaries of the puncta. We then algorithmically identify the enclosed regions formed by the boundaries and counted the number of puncta in each figure. For the case of co-localisation, the co-ordinates of the centres of the punctas/clusters from each channel were calculated and compared to see if they lie within a small tolerance for each puncta and channel. The puncta/clusters satisfying the abovementioned conditions were considered to be co-localised and were counted.

