## Supplementary figures for "The Irish potato famine pathogen subverts host vesicle trafficking to channel starvation-induced autophagy to the pathogen interface"

**Supplementary materials:**

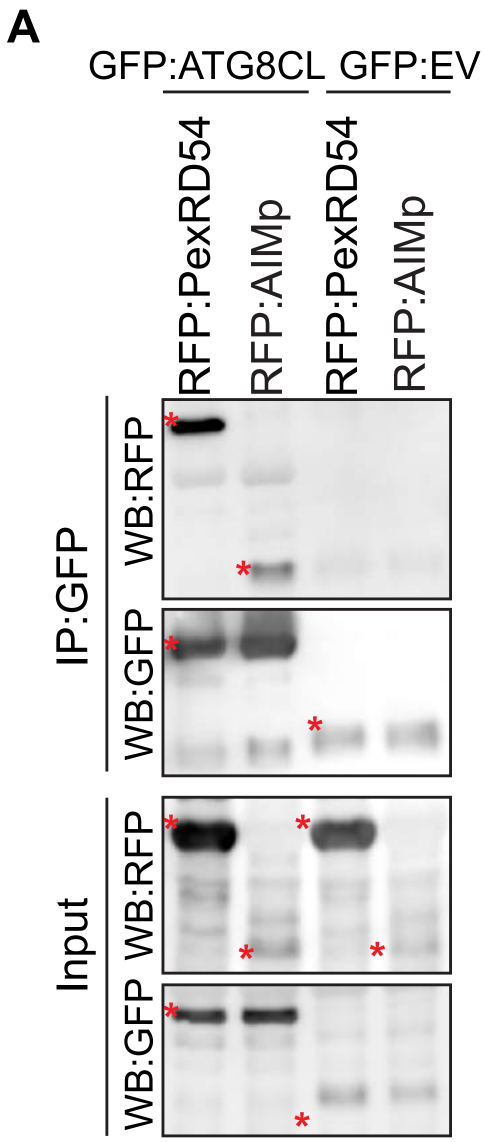


**Figure S1. PexRD54 and AIMp interact with ATG8CL *in vivo.*** *In planta* co-immunoprecipitation between ATG8CL and PexRD54 or AIMp. GFP:ATG8CL was transiently co-expressed with either RFP:PexRD54 or RFP:AIMp. IPs were obtained with anti-GFP antiserum. Red asterisks indicate expected band sizes.

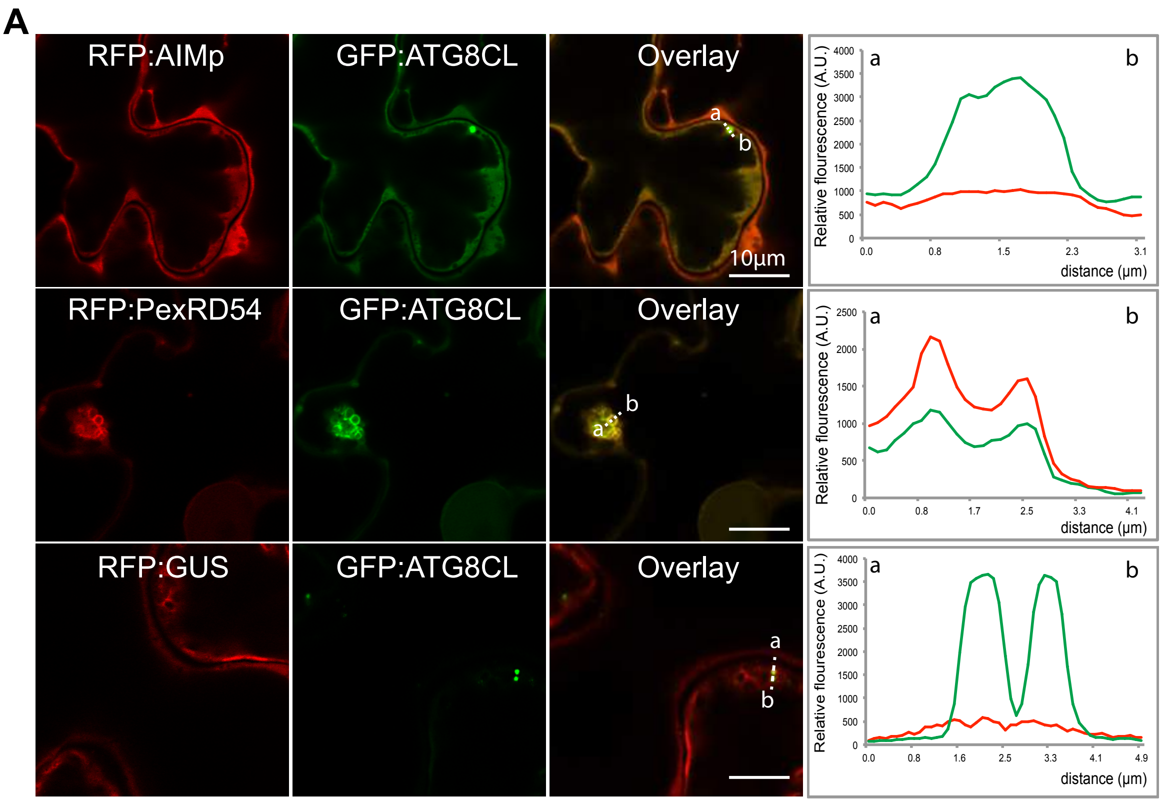


**Figure S2. PexRD54 co-localizes with ATG8CL puncta.** Maximum projection confocal micrographs of *N. benthamiana* leaf epidermal cells transiently expressing either RFP:AIMp (top), RFP:PexRD54 (middle), or RFP:GUS (bottom), with GFP:ATG8CL. Transects in overlay panel correspond to plot of relative fluorescence over the labelled distance. RFP:PexRD54 co-localises in discrete puncta with GFP:ATG8CL while RFP:AIMp and RFP:GUS show diffuse expression through GFP:ATG8CL puncta. Images shown are maximal projections of 12 frames with 1 µm steps. Scale bars, 10 µm.

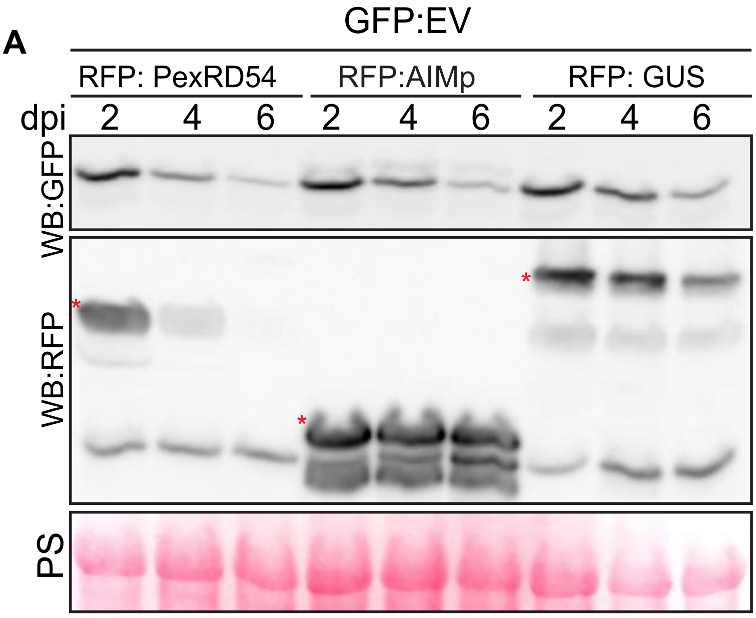


**Figure S3. AIMp does not stabilize EV:GFP.** Western blotting shows EV:GFP is not stabilized by co-expression with RFP:PexRD54, RFP:AIMp, or RFP:GUS. Total protein extracts were isolated 2, 4 and 6 days after infiltration and immunoblotted.

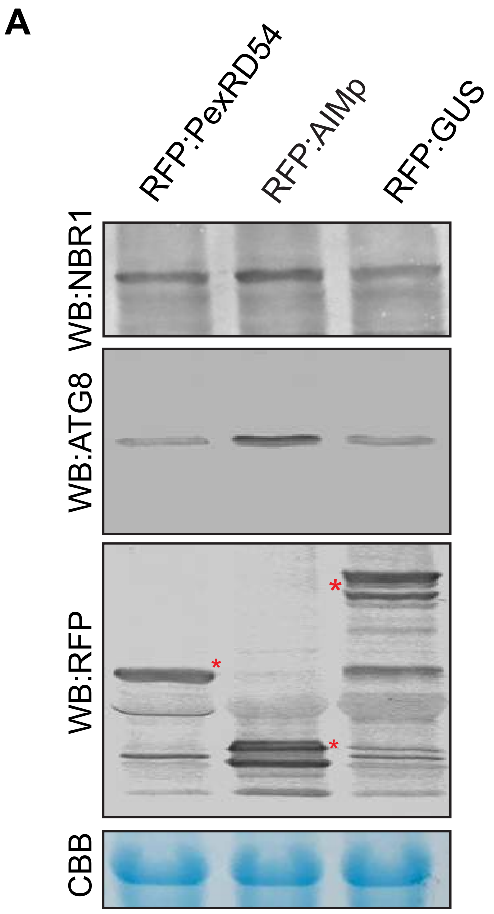


**Figure S4. AIMp stabilizes endogenous NBR1/Joka2 and ATG8(s).** RFP:AIMp stabilizes endogenous NBR1/Joka2 and ATG8(s). PexRD54 has a milder effect on NBR1/Joka2 stabilization but no apparent effect on ATG8(s). Total protein extracts were isolated 3 days after infiltration and immunoblotted.

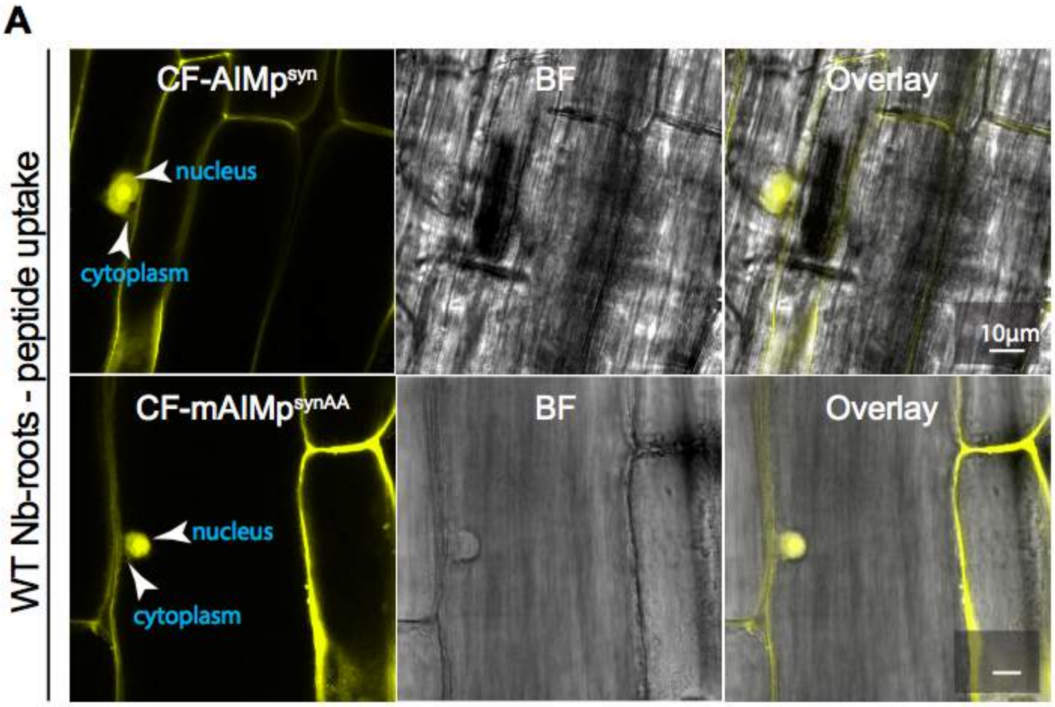


**Figure S5. Cell penetrating AIM peptide constructs are uptaken in roots.** Maximum projection confocal micrographs of *N. benthamiana* root epidermal cells exogenously supplied with either CF-AIMp$^{syn}$ (top) or CF-mAIMp$^{syn}$ (bottom). Both AIMp and its mutant form are translocated well by the roots into the cytoplasm and diffuse into the nucleus. CF = 5-Carboxyfluorescein. Images shown are maximal projections of 10 frames with 1 µm steps. Scale bars, 10µm.

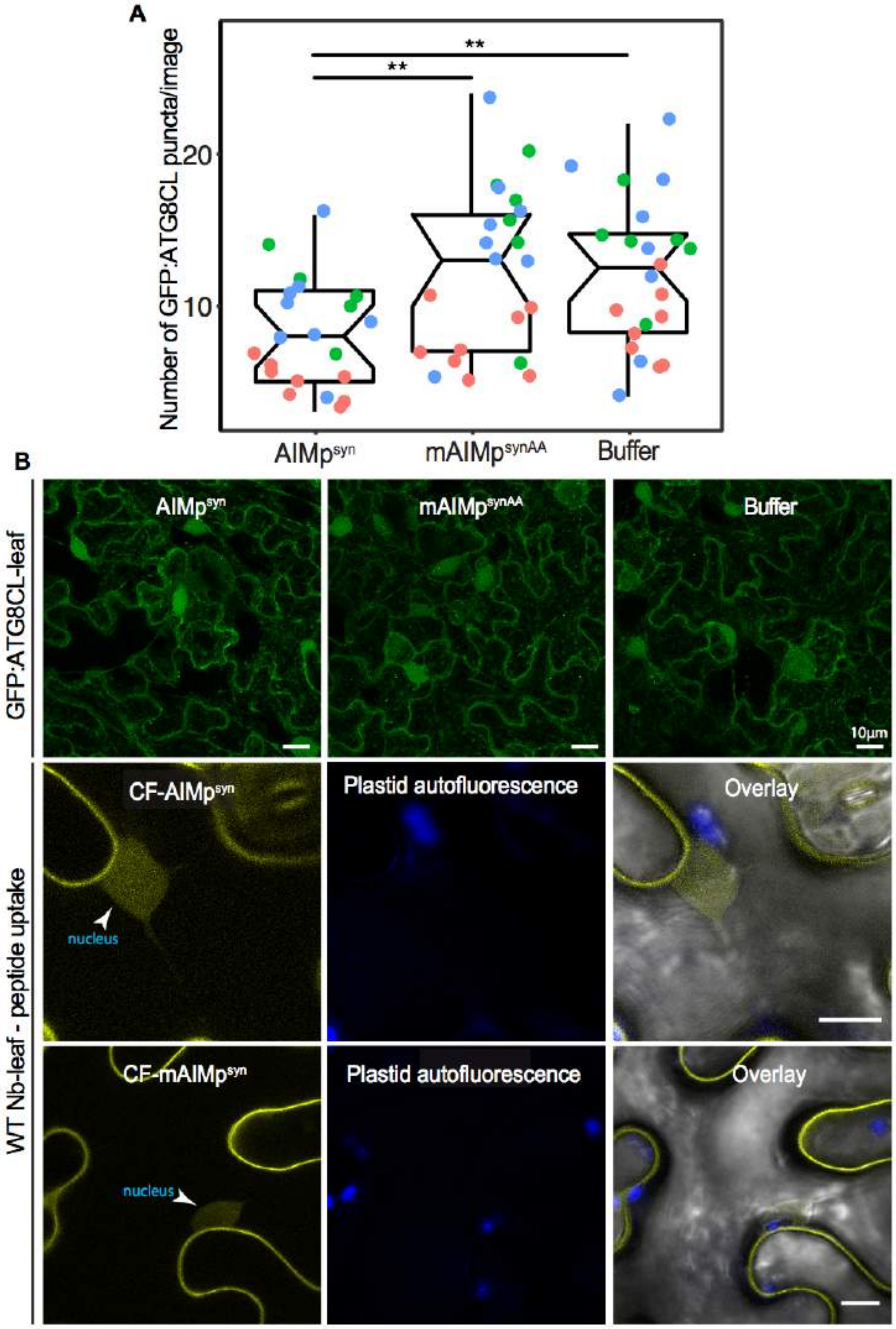


**Figure S6. Cell penetrating AIM peptides translocate inside leaf cells and result in decreased numbers of ATG8CL puncta.** (A) Scatter-boxplot shows exogenous application of cell penetrating AIMp$^{syn}$ in GFP-ATG8CL transgenic *N. benthamiana* significantly decreases the number of ATG8 puncta per image (8, *N* = 21 images quantified) compared to cell penetrating mAIM$^{synAA}$ (12, *N* = 22 images quantified), or Buffer control (12, *N* = 22 images quantified). Scattered points show individual data points, color indicates biological repeat. Representative maximum projection confocal micrographs of epidermal cells. (B) Single-plane confocal micrographs of WT *N. benthamiana* exogenously supplied with 5-Carboxyfluorescein (CF) tagged versions of AIM$^{syn}$ and mAIM$^{syn}$, showing peptide translocation into the nucelocytoplasm of leaf epidermal cells. Images shown are maximal projections of 25 frames with 1 µm steps. Scale bars, 10µm.

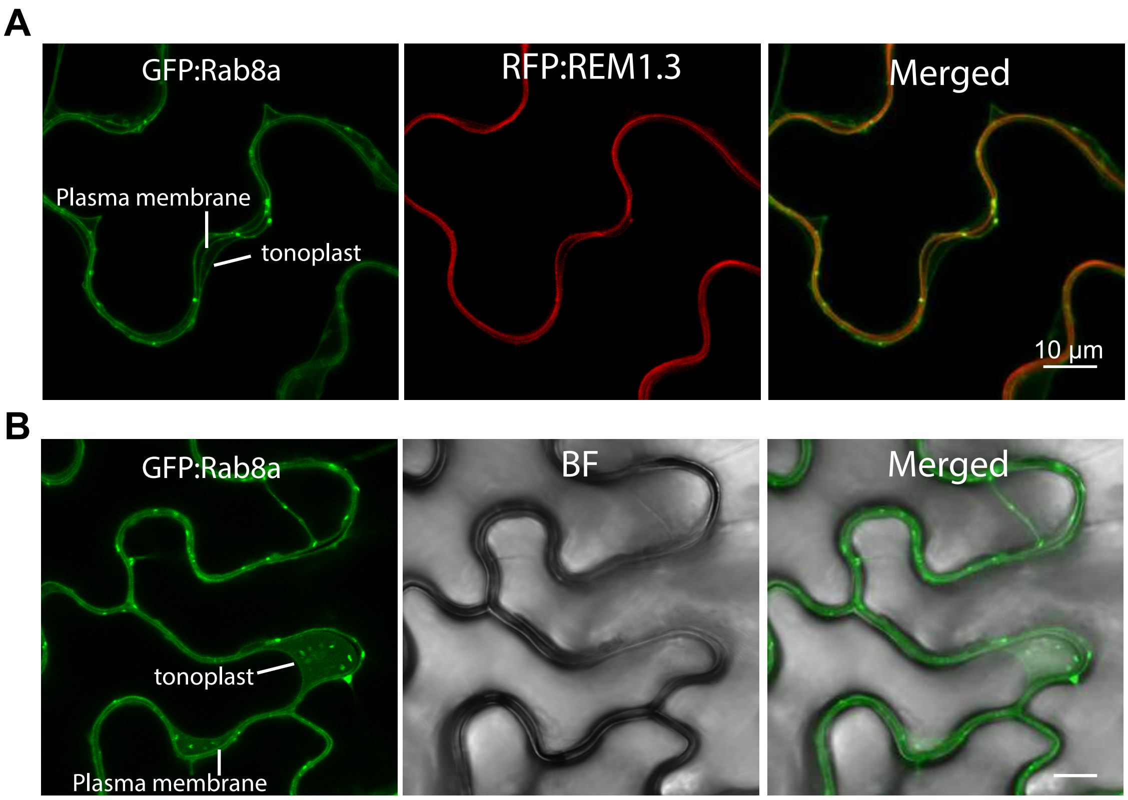


**Figure S7. Rab8a localizes in the tonoplast, plasma membrane and puncta.** Single-plane confocal micrographs of *N. benthamiana* leaf epidermal cells transiently expressing either GFP:Rab8a alone (top) or GFP:Rab8a with the plasma membrane marker RFP:REM1.3 (bottom). Rab8a localises in discrete puncta as well as in the plasma membrane with RFP:REM1.3 and in the tonoplast. Images shown are maximal projections of 15 frames with 1 μm steps. Scale bars, 10 μm.

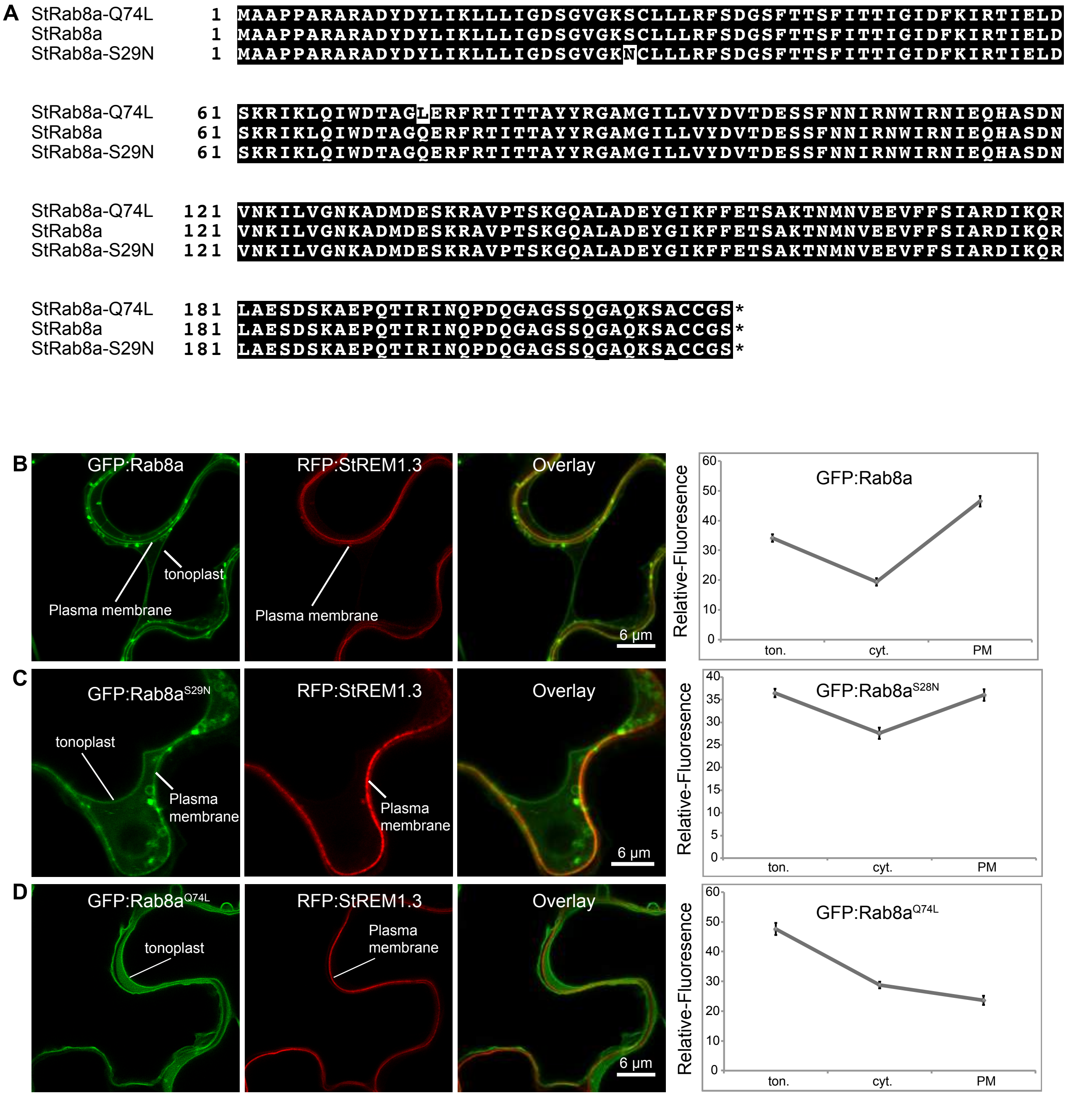


**Figure S8. Subcellular distribution of Rab8a and its mutants**. (A) Amino acid sequences of *S. tuberosum* Rab8a$^{Q74L}$ (top), wild type Rab8a (middle) and Rab8a$^{S29N}$ (bottom) proteins. (B) Maximum projection confocal micrographs of *N. benthamiana* leaf epidermal cells transiently expressing GFP:Rab8a (top), Rab8a$^{S29N}$ (middle), and Rab8a$^{Q74L}$ (bottom), with plasma membrane marker RFP:REM1.3. Transects in overlay panel correspond to plot of relative fluorescence over the labelled distance. Images shown are maximal projections of 20 frames with 1 µm steps. Scale bars, 6 µm.

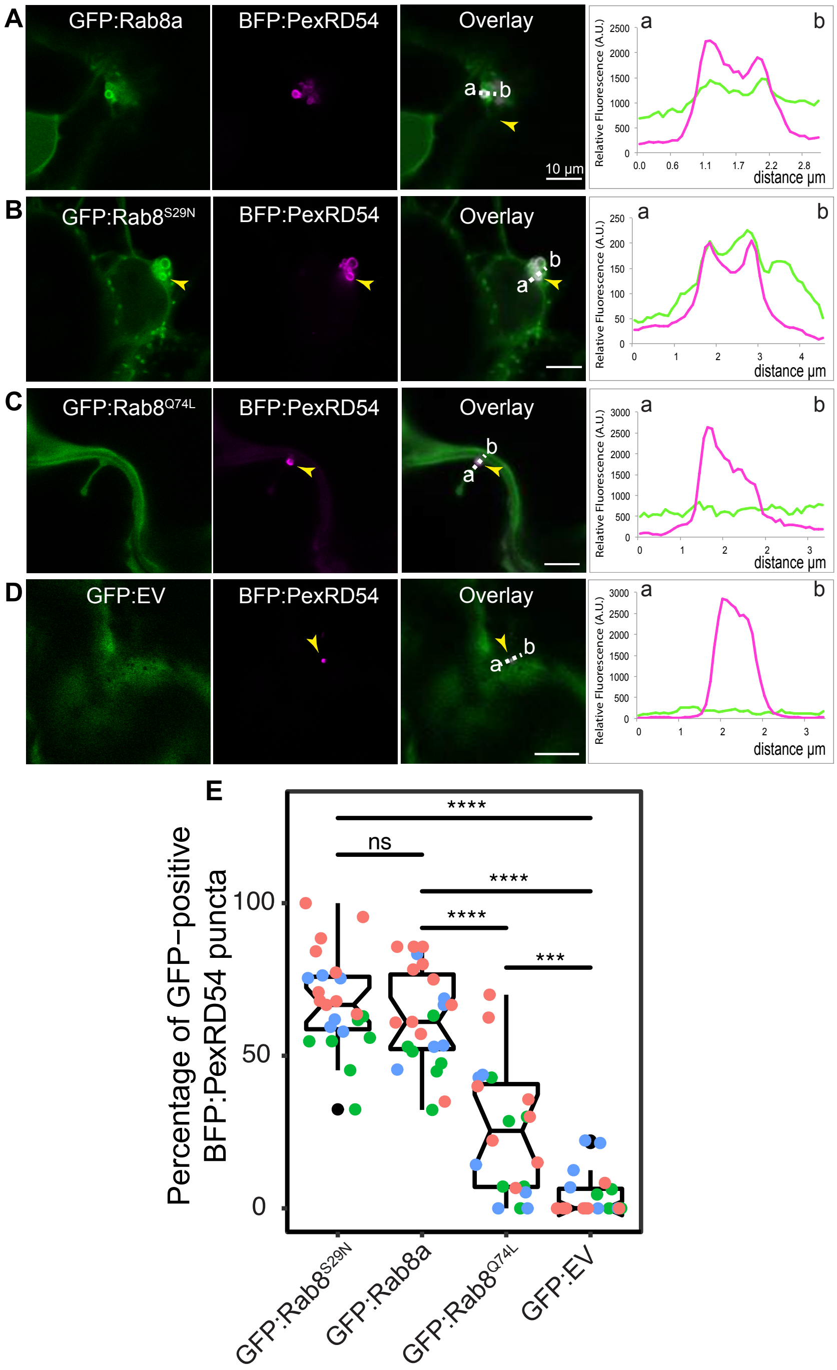


**Figure S9. PexRD54 puncta preferentially co-localize with puncta of wild type and GDP bound form of Rab8a (Rab8a$^{S29N}$) rather than GTP bound form (Rab8a$^{Q74L}$).** Maximum projection confocal micrographs of *N. benthamiana* leaf epidermal cells transiently expressing BFP:PexRD54 with GFP:Rab8a (A), GFP:Rab8a$^{S29N}$ (B), GFP:Rab8a$^{Q74L}$ (C), or GFP:EV (D). Transects in overlay panel correspond to plot of relative fluorescence over the labelled distance. GFP:Rab8a and GFP:Rab8a$^{S29N}$ shows similar co-localization pattern with BFP:PexRD54, while GFP:Rab8a$^{Q74L}$ has much lower co-localization with BFP:PexRD54. Images shown are maximal projections of 15 frames with 1 μm steps Scale bars, 10μm (E) Scatter-boxplot shows GFP:Rab8a$^{Q74L}$ (25%, N = 20 images quantified) colocalizes significantly less with PexRD54 compared to GFP:Rab8a (68%, N = 23 images quantified), GFP:Rab8a$^{S29N}$ (67%, N = 23 images quantified) or an empty vector (11%, N = 23 images quantified).

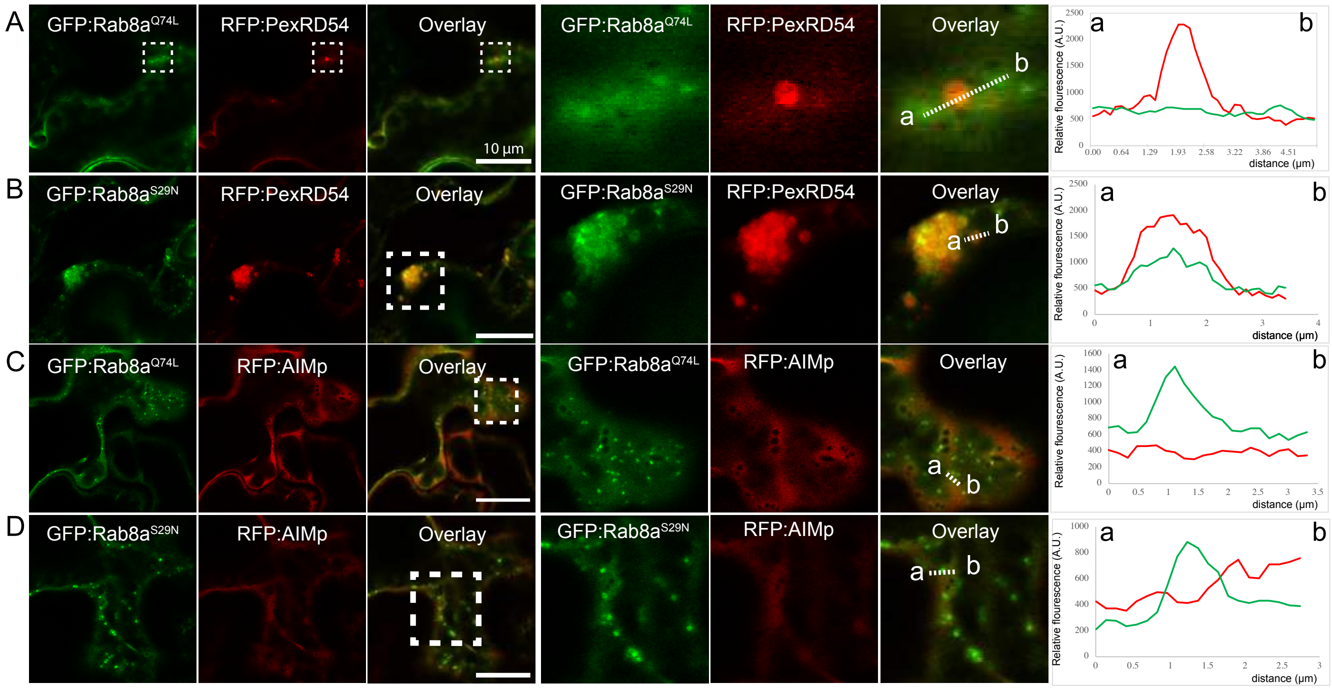


**Figure S10. Rab8a$^{S29N}$ and GFP:Rab8a$^{Q74L}$ do not co-localize with AIM peptide *in planta*.** Maximum projection confocal micrographs of *N. benthamiana* leaf epidermal cells transiently expressing either RFP:PexRD54 or RFP:AIMp, with either (B, D) GFP:Rab8a$^{S29N}$ or (A, C) GFP:Rab8a$^{Q74L}$. While GFP:Rab8a$^{S29N}$, GDP bound form, labels PexRD54 puncta, GDP bound form of Rab8a, GFP:Rab8a$^{Q74L}$, does not label most of the RFP:PexRD54 puncta. Both of the mutants do not co-localize with RFP:AIMp. Confocal micrographs on the right are zoomed versions of white dashed squares on the left images. Transects in overlay panel correspond to plot of relative fluorescence over the labelled distance. Images shown are maximal projections of 21 frames with 1 µm steps. Scale bars, 10 µm.

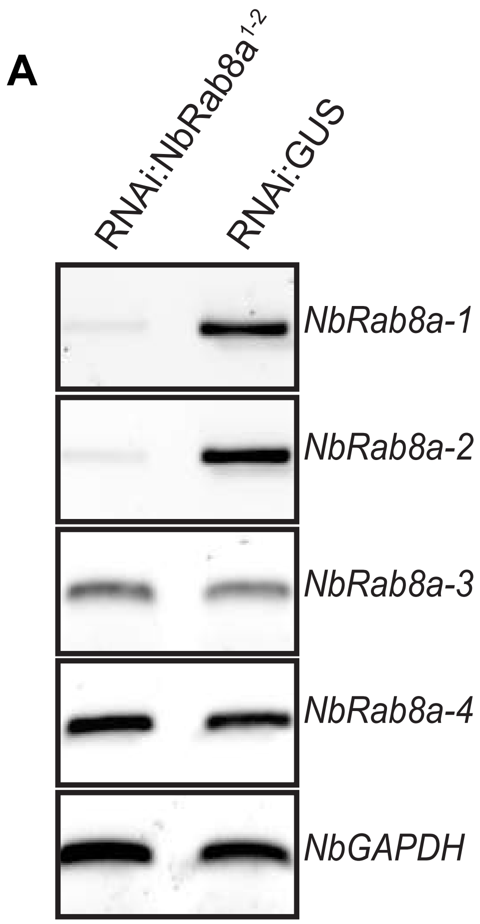


**Figure S11. Validation of *NbRab8a-1* and *NbRab8a-2* silencing by RNAi:NbRab8a$^{1\text{-}2}$.** Constructs carrying hairpin plasmids (pRNAi-GG) targeting *NbRab8a,* or *GUS* reporter gene were infiltrated to *N. benthamiana* and the expression of targeted genes was assessed by RT-PCR at three days post silencing. RT-PCR verified efficient gene silencing of *NbRab8a-1* and *NbRab8a-2.* Glyceraldehyde 3-phosphate dehydrogenase (GAPDH) was used as internal control. cDNA was synthesized using total RNA.

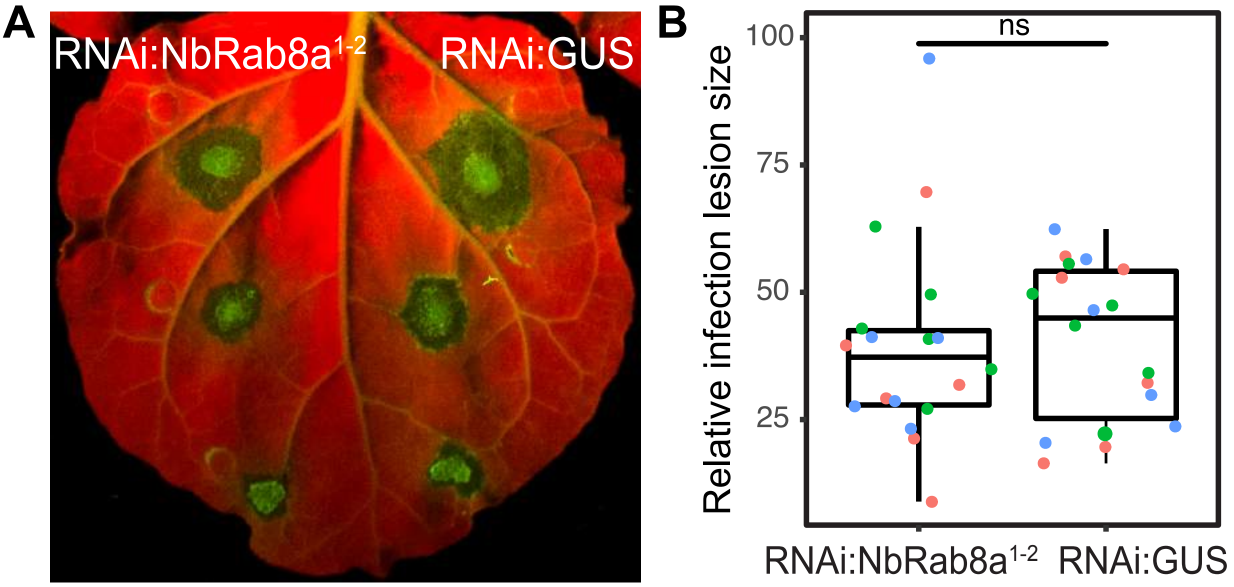


**Figure S12. Silencing *Rab8a1-2* does not affect susceptibility to *P. infestans*.** (A-B) Silencing of homologs 1-2 of Rab8a (41, *N* = 17 infected leaves) does not significantly affect *P. infestans* necrotic lesion size compared to a silencing control (39, N = 19 infected leaves). *N. benthamiana* leaves expressing RNAi:NbRab8a$^{1\text{-}2}$ or RNAi:GUS were infected with *P. infestans* and pathogen growth was determined by measuring infection lesion size 7 days post-inoculation.

**A**

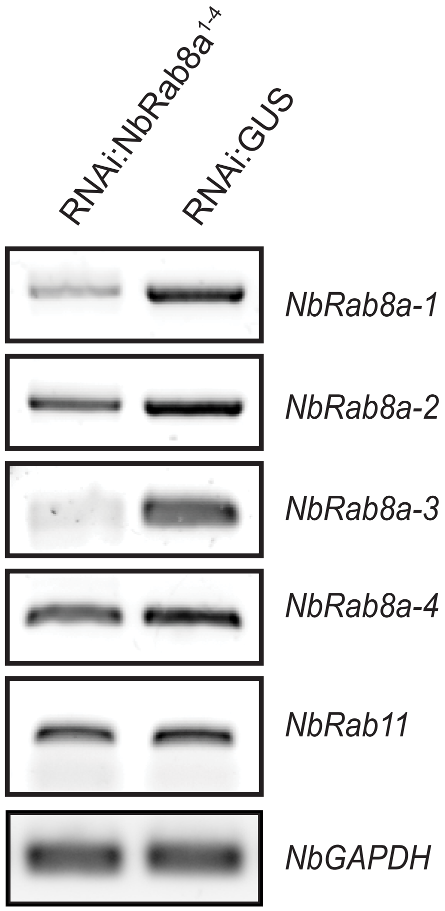


**Figure S13. Validation of RNAi:NbRab8a$^{1-4}$ silencing construct.** RT-PCR validates efficient silencing of *NbRab8a1-4* when RNAi:NbRab8a$^{1-4}$ is expressed, compared to RNAi:GUS expression. Rab11 is not efficiently silenced by RNAi:NbRab8a$^{1-4}$ compared to RNAi:GUS.

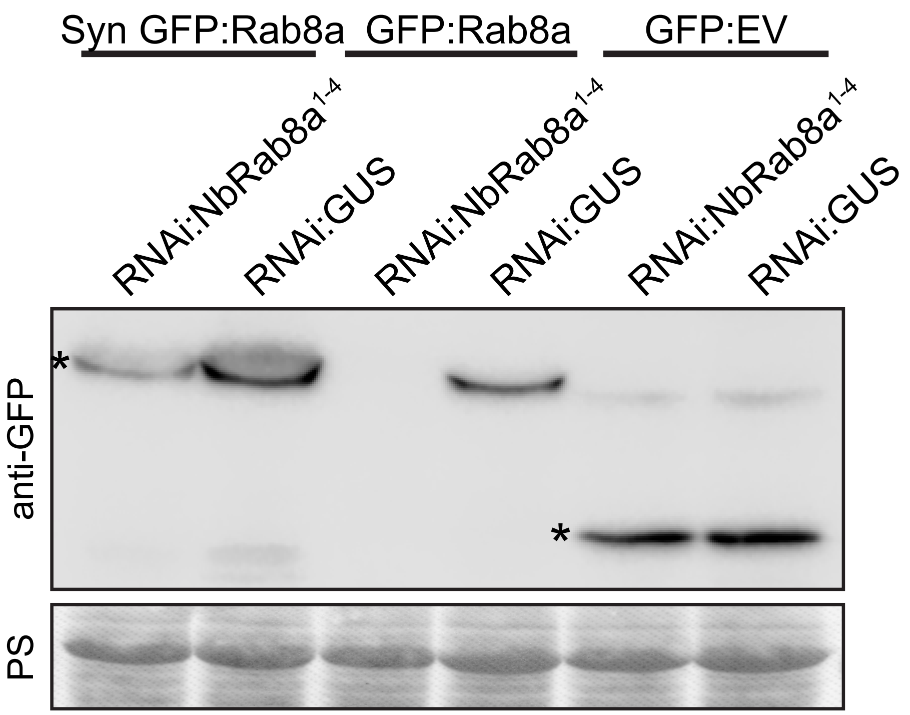


**Figure S14. Synthetic GFP:Rab8a is more resistant to silencing by RNAi:NbRab8a$^{1-4}$ compared to GFP:Rab8a.** Western-blot analysis shows that Syn GFP:Rab8a shows significantly higher protein levels compared to GFP:Rab8a when RNAi:NbRab8a$^{1-4}$ is co-expressed. RNAi:NbRab8a$^{1-4}$ does not affect GFP:EV protein levels compared to RNAi:GUS. Total protein extracts were isolated 4 days after infiltration and immunoblotted.

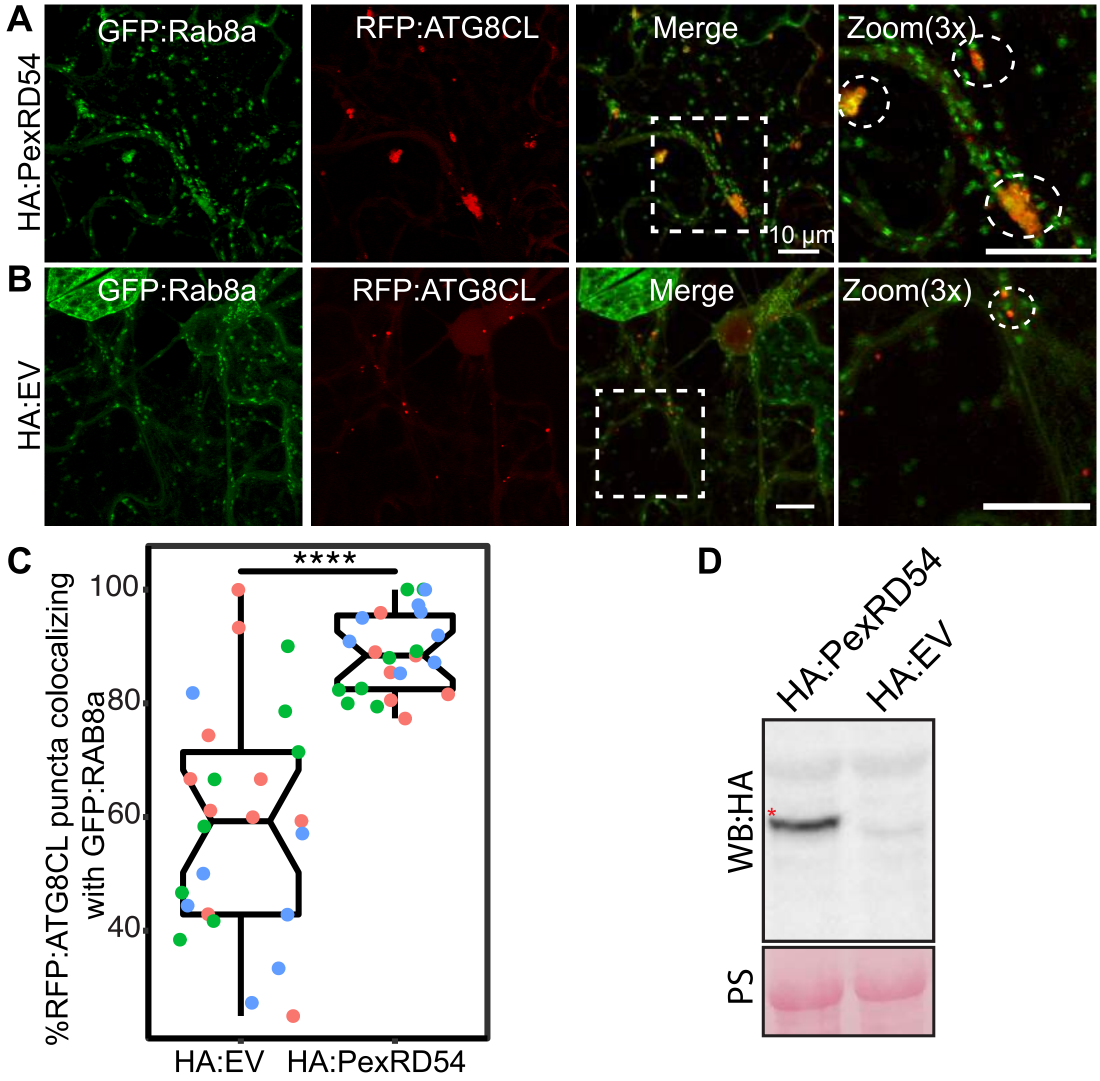


**Figure S15. PexRD54 increases Rab8a and ATG8CL co-localisation.** (A-B) Maximum projection confocal micrographs of *GFP:NbRab8a* transgenic *N. benthamiana* leaf epidermal cells transiently expressing either (A) HA:PexRD54, or (B) HA:EV, with GFP:ATG8CL and RFP:ATG8CL. Clustering of ATG8CL with Rab8a increases in the presence of PexRD54 expression. Images shown are maximal projections of 22 frames with 1 µm steps, scale bars 10um. (C) Scatter-boxplot shows HA:PexRD54 expression significantly increases the percentage of RFP:ATG8CL puncta colocalising with GFP:Rab8a per image (87%, *N* = 22 images quantified) compared to expressing HA:EV control (58%, *N* = 25 images quantified). (D) Western blot analysis showing HA:PexRD54 is well expressed and the expected size. Total protein extracts were isolated 3 days after infiltration and immunoblotted.

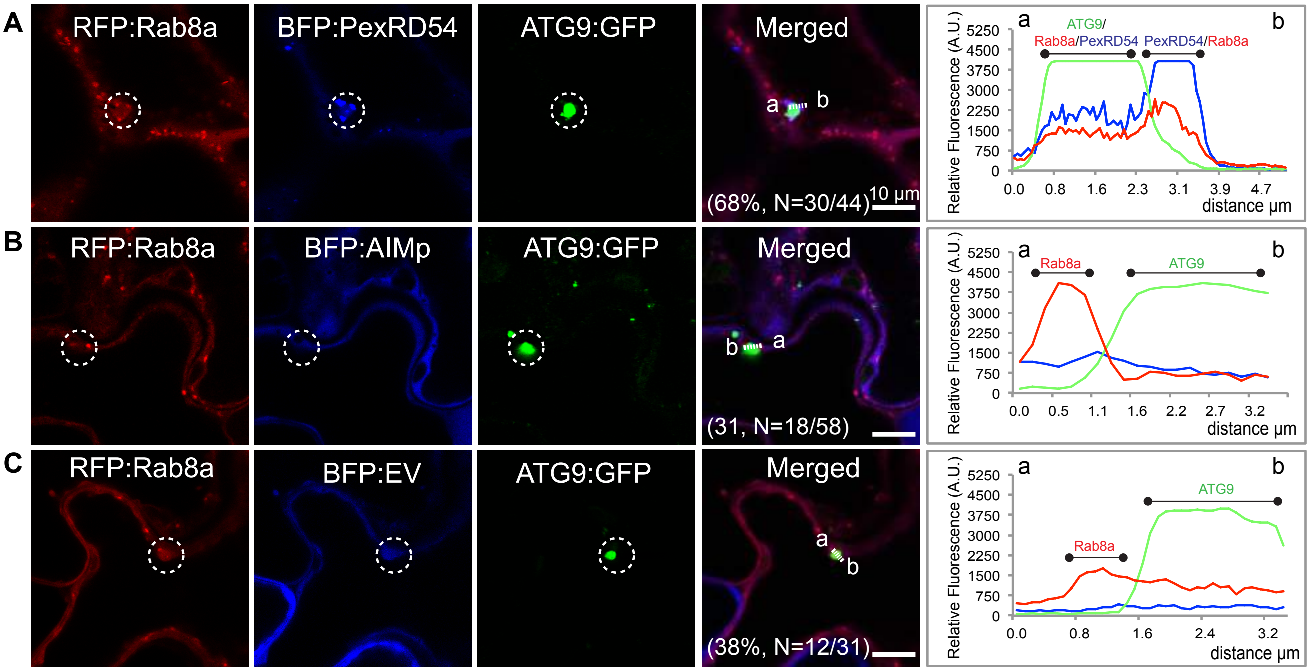


**Figure S16. PexRD54 enhances Rab8a/ATG9 colocalisation.** Maximum projection confocal micrographs (xyz) of *N. benthamiana* leaf epidermal cells transiently expressing RFP:Rab8a and ATG9:GFP with either (A) BFP:PexRD54, (B) BFP:AIMp or (C) or BFP:EV. Transects in overlay panel correspond to plot of relative fluorescence over the labelled distance. Images shown are maximal projections of 15 frames with 1 µm steps. Scale bars, 10 µm. Expression of BFP:PexRD54 increases the amount of ATG9 positive Rab8a puncta (68%, $N$ = 44 images quantified) compared to a BFP control (38%, $N$ = 31 images quantified). Expression of BFP:AIMp (31%, $N$ = 58 images quantified) does not have an effect compared to a BFP control.

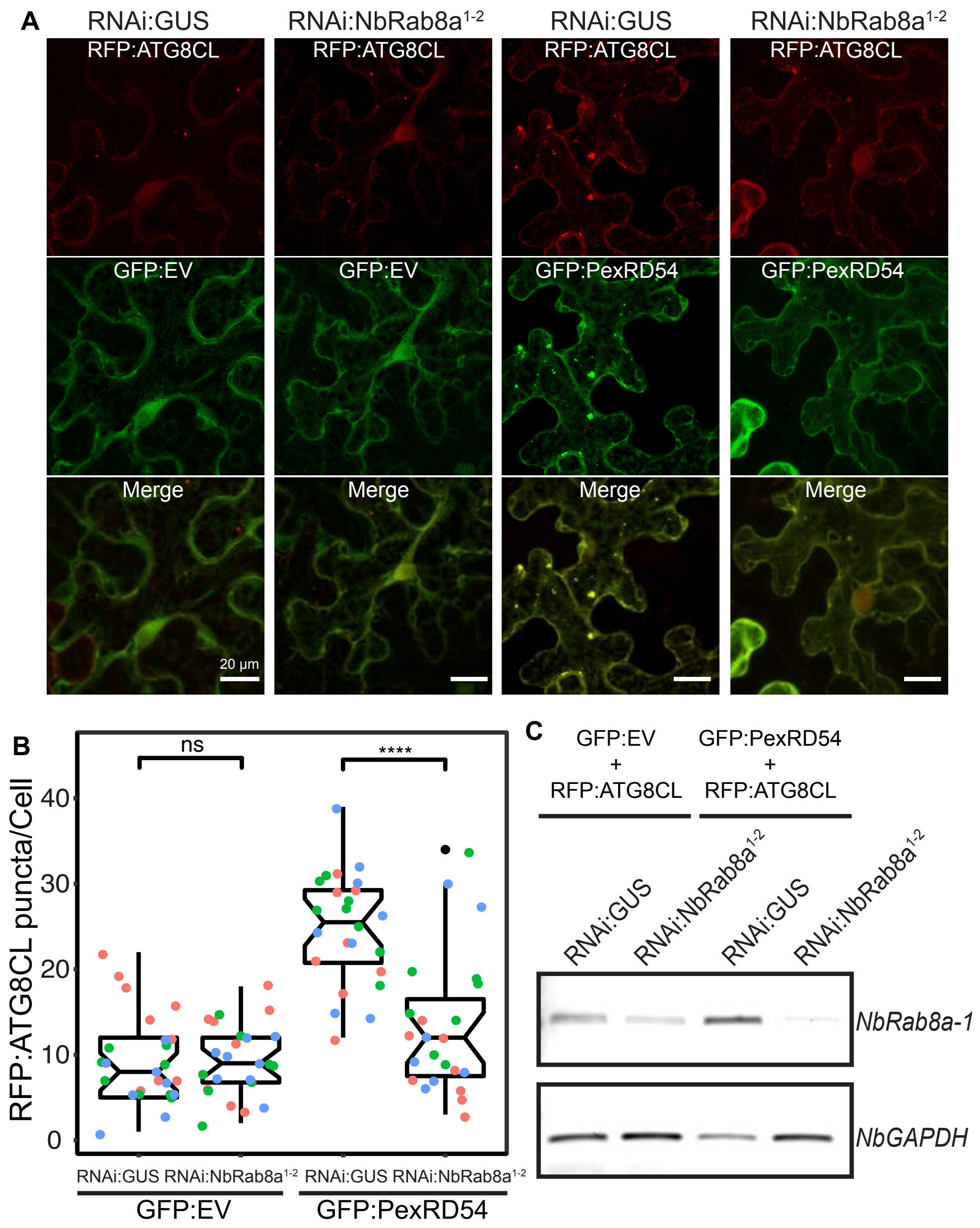


**Figure S17. PexRD54 increases the number of ATG8CL puncta in a Rab8a-dependent manner.** (A) Maximum projection confocal micrographs of *N. benthamiana* leaf epidermal cells transiently expressing RFP:ATG8CL and GFP:EV with either RNAi:GUS (left column) or RNAi:NbRab8a$^{1\text{-}2}$ (right column). RFP:ATG8CL puncta are still seen in Rab8a-silenced tissue. Maximum projection confocal micrographs of *N. benthamiana* leaf epidermal cells transiently expressing RFP:ATG8CL and GFP:PexRD54 with either RNAi:GUS (left column) or RNAi:NbRab8a$^{1\text{-}2}$ (right column). RFP:ATG8CL puncta are reduced in Rab8a-silenced tissue. Images shown are maximal projections of 22 frames with 1 µm steps. Scale bars, 10um. (B) Scatter-boxplot shows quantification of A-B. When GFP:PexRD54 is co-expressed, RNAi:NbRab8a$^{1\text{-}2}$ expression significantly reduces the number of RFP:ATG8CL puncta per cell (13, *N* = 23 images quantified) compared to RNAi:GUS expression (25, *N* = 24 images quantified), but a similar reduction is not seen when GFP:EV is co-expressed(9, N = 24; 9, N = 25 images quantified). (C) RT-PCR validates efficient silencing of *NbRab8a-1* in experiment A-C when RNAi:NbRab8a$^{1\text{-}2}$ is expressed, compared to RNAi:GUS expression.

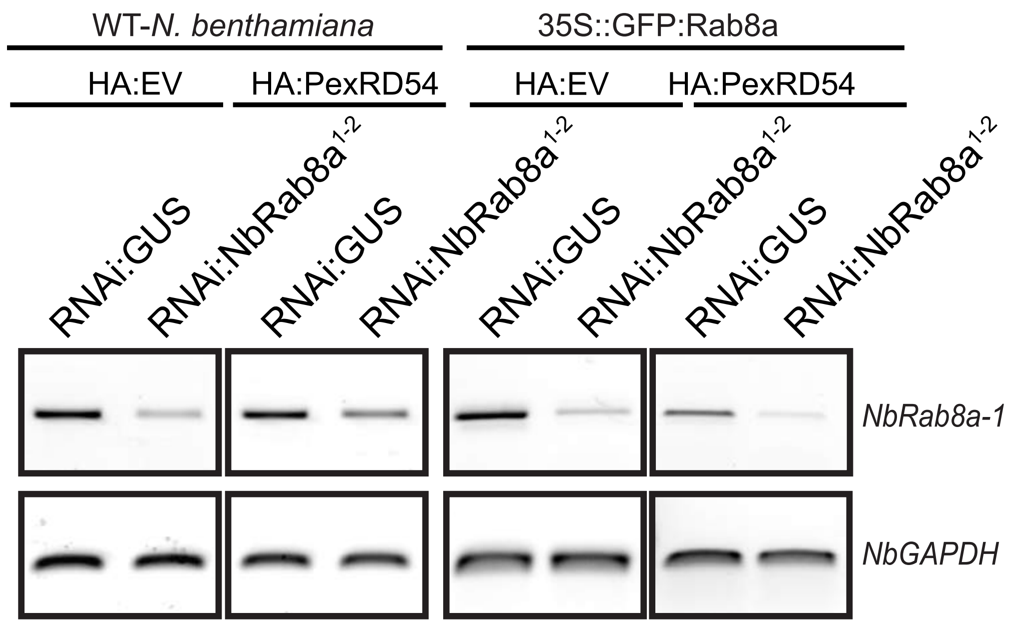


**Figure S18. Validation of silencing of *NbRab8a-1* by RNAi:NbRab8a$^{1\text{-}2}$.** RT-PCR validates silencing of *NbRab8a-1* for Fig5A-B when RNAi:NbRab8a$^{1\text{-}2}$ is expressed, compared to RNAi:GUS expression

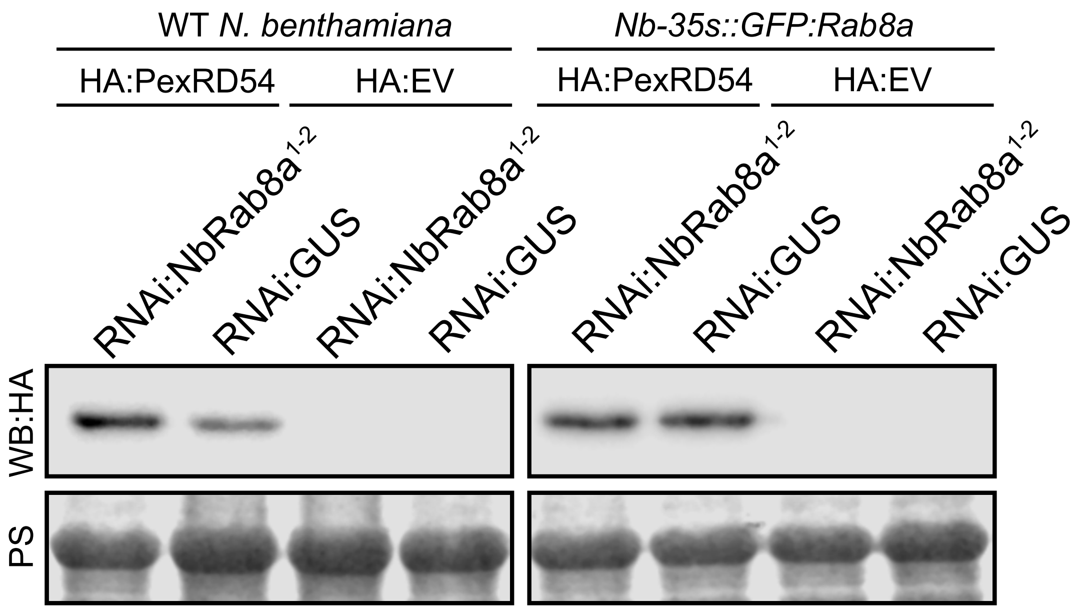


**Figure S19. Silencing of *NbRab8a1-2* does not reduce PexRD54 protein levels.** Expression of RNAi:NbRab8a$^{1-2}$ does not reduce PexRD54 protein levels. Total protein extracts were isolated 3 days after infiltration and immunoblotted.

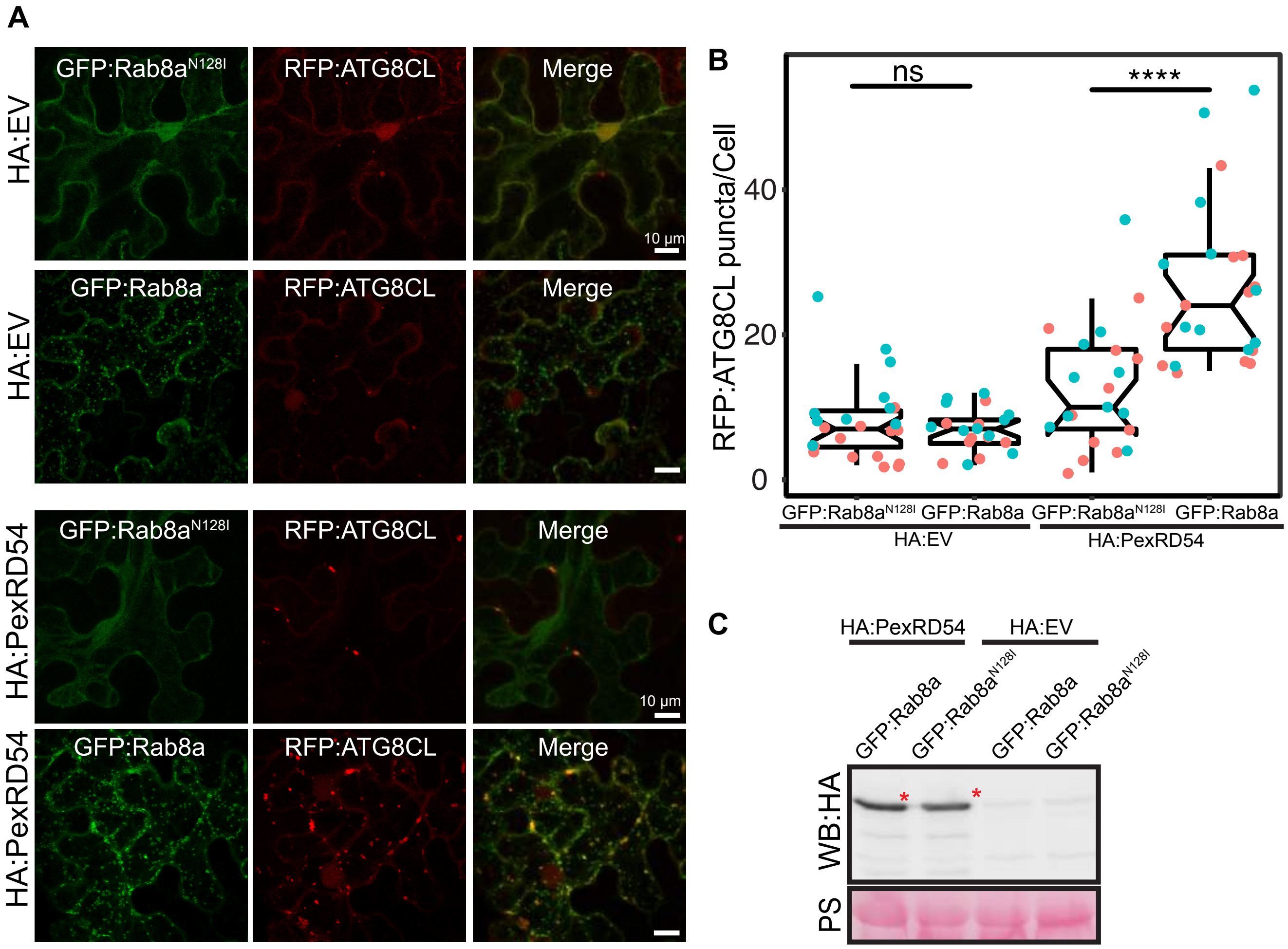


**Figure S20. Dominant negative mutant of Rab8a (N128I) decreases PexRD54 induced autophagosome formation.** (A) Maximum projection confocal micrographs of *N. benthamiana* leaf epidermal cells transiently expressing RFP:ATG8CL with either GFP:Rab8a$^{N128I}$ (top) or GFP:Rab8a (bottom) together with either HA:EV or HA:PexRD54. RFP:ATG8CL puncta are still seen in Rab8a-silenced tissue. GFP:Rab8a$^{N128I}$ significantly reduces autophagosome number induced by HA:PexRD54. Images shown are maximal projections of 22 frames with 1 µm steps Scale bars, 10 µm. (B) Quantification of RFP:ATG8CL puncta is shown on scatter-boxplot. Expression of GFP:Rab8a$^{N128I}$ significantly reduces the amount of HA:PexRD54 triggered RFP:ATG8CL autophagosomes (12, *N* = 20 images quantified) compared to GFP:Rab8a (26, *N* = 23 images quantified). When HA:EV is expressed, expression of GFP:Rab8a$^{N128I}$ does not affect the amount of RFP:ATG8CL autophagosomes (8, *N* = 23 images quantified) compared to GFP:Rab8a (8, *N* = 23 images quantified). (C) Western blot analysis showing HA:PexRD54 is expressed at similar levels when co-expressed with GFP:Rab8a$^{N128I}$ or GFP:Rab8a.

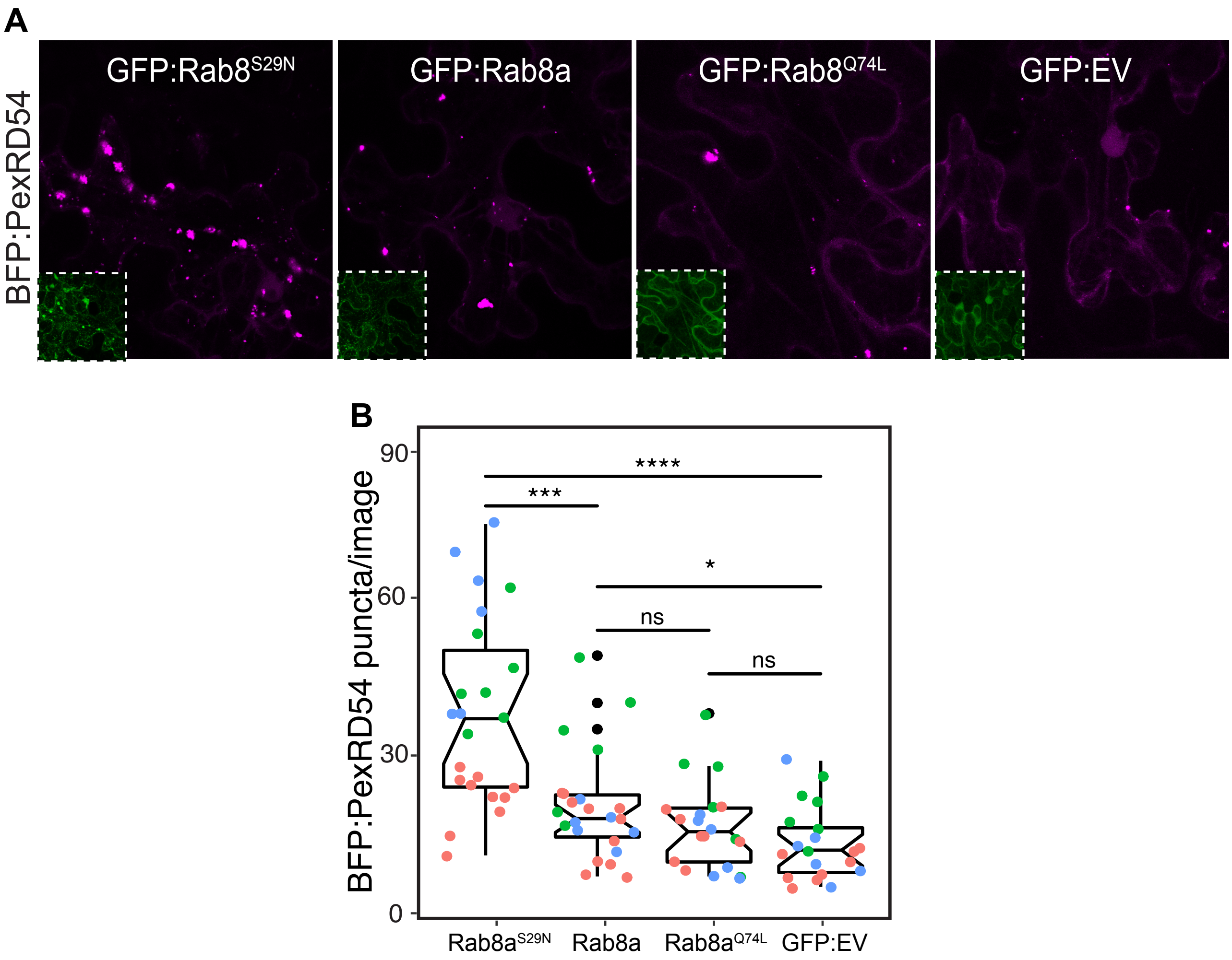


**Figure S21. WT Rab8a and Rab8a$^{S29N}$ increases the amount of PexRD54 triggered puncta.** (A) Confocal micrographs of *N. benthamiana* leaf epidermal cells transiently expressing BFP:PexRD54 with either GFP:Rab8a$^{S29N}$, GFP:Rab8a, GFP:Rab8a$^{Q74L}$, or GFP:EV. (B) Overexpression of GFP:Rab8a$^{S29N}$ enhances PexRD54 puncta number (38, $N$ = 23 images quantified) compared to the GTP mutant form GFP:Rab8a$^{Q74L}$ (17, $N$ = 20 images quantified) or an empty vector control (13, $N$ = 20 images quantified). The WT Rab8a had a mildly positive effect on PexRD54 puncta number (20, $N$ = 23 images quantified) compared to an empty vector. (B, D, F) Scattered points show individual data points, colours indicate biological repeats. Scale bars, 10 µm.

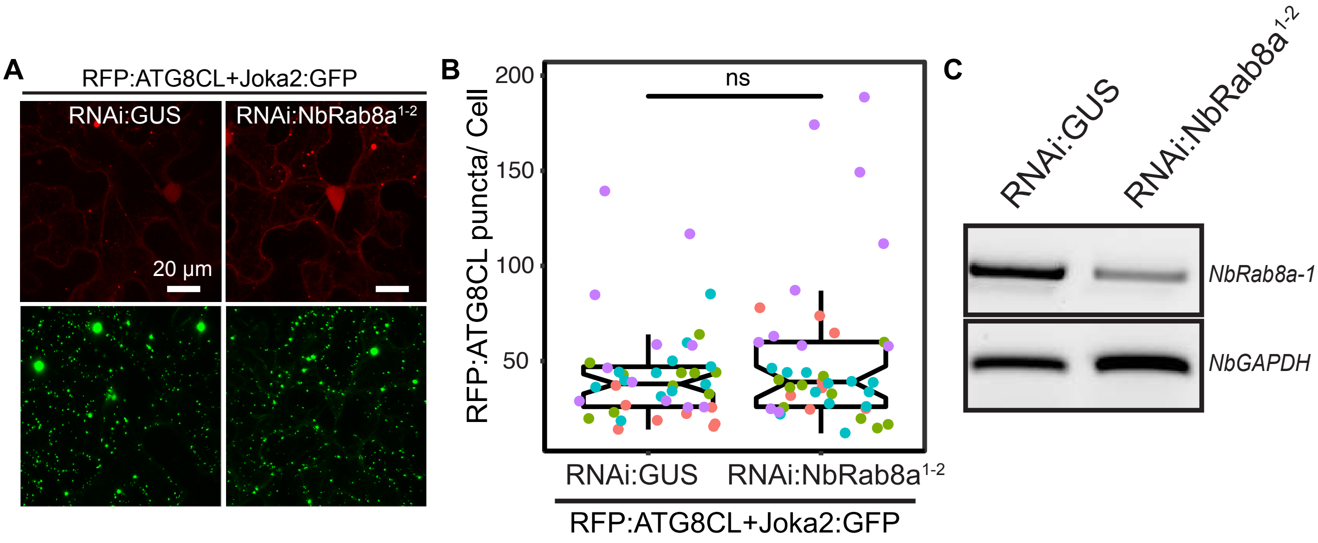


**Figure S22. Expression of RNAi:NbRab8a$^{1\text{-}2}$ in *N. benthamiana* does not affect Joka2-induced autophagosome formation**. (A) Maximum projection confocal micrographs of *N. benthamiana* leaf epidermal cells transiently expressing RFP:ATG8CL and Joka2:GFP, with either RNAi:GUS or RNAi:Rab8a$^{1\text{-}2}$. Scale bars, 10µm Images shown are maximal projections of 23 frames with 1 µm steps (B) Scatter-boxplot shows that silencing *NbRab8a1-2* (51, *N* = 41 images quantified) does not affect induction of ATG8CL autophagosome formation by Joka2 compared to a GUS silencing control (42, *N* = 42 images quantified). (C) RT-PCR verified gene silencing of *NbRab8a-1.* Glyceraldehyde 3-phosphate dehydrogenase (GAPDH) was used as internal control.

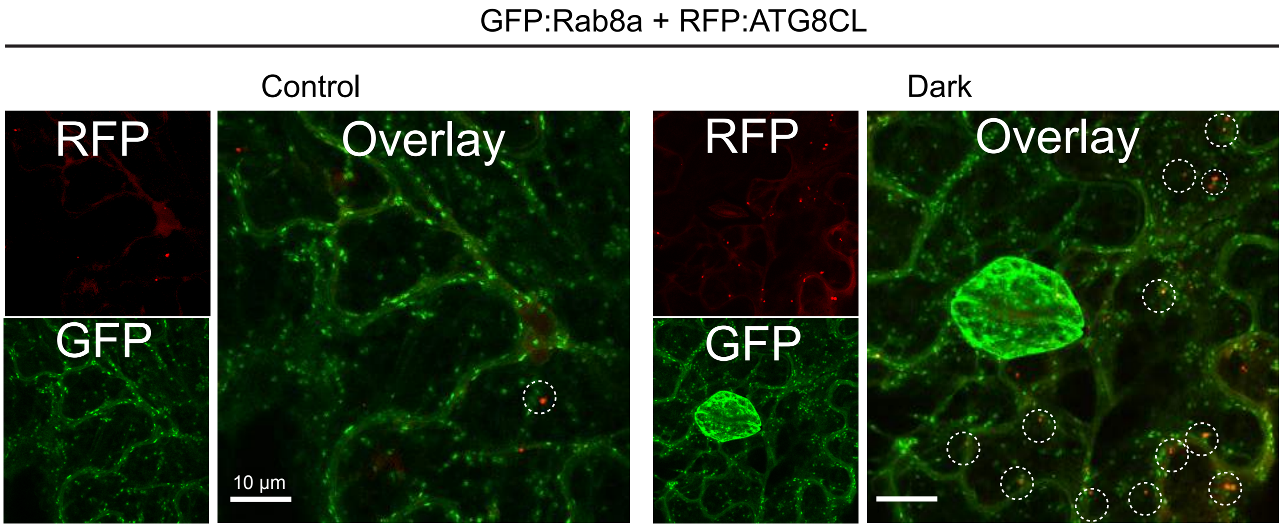


**Figure S23. Dark treatment increases ATG8CL/Rab8a colocalization.** Maximum projection confocal micrographs of Nb::35SGFP:Rab8a *N. benthamiana* leaf epidermal cells transiently expressing RFP:ATG8CL under normal light or 24-hour-dark conditions.

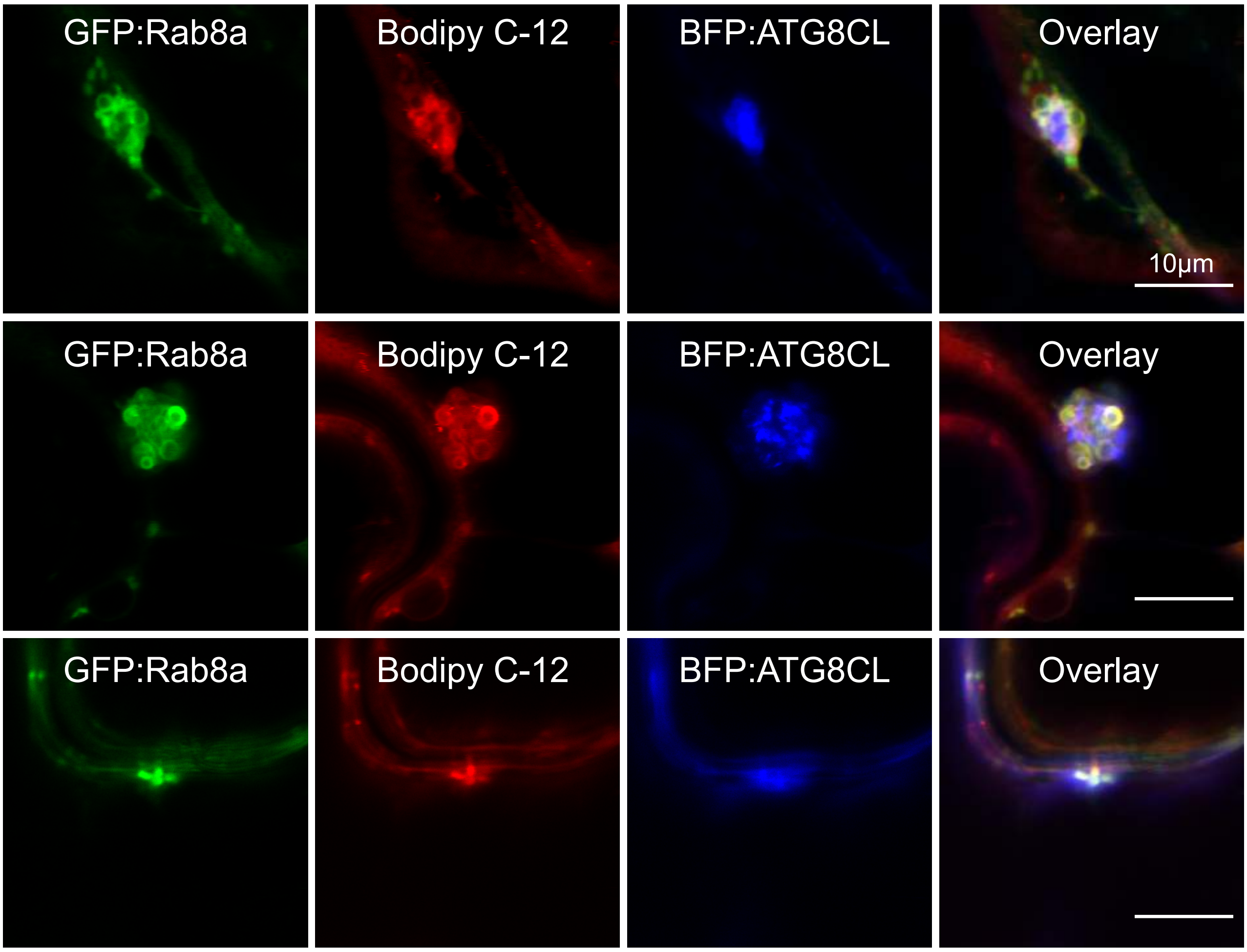


**Figure S24. Rab8a colocalises in puncta and ring-shaped vesicle-like structures with ATG8CL and Bodipy C-12.** Maximum projection confocal micrographs of *N. benthamiana* leaf epidermal cells transiently expressing GFP:Rab8a, BFP:ATG8CL and stained with Bodipy C-12. Images shown are maximal projections of 15 frames with 1 µm steps. Scale bars, 10 µm.

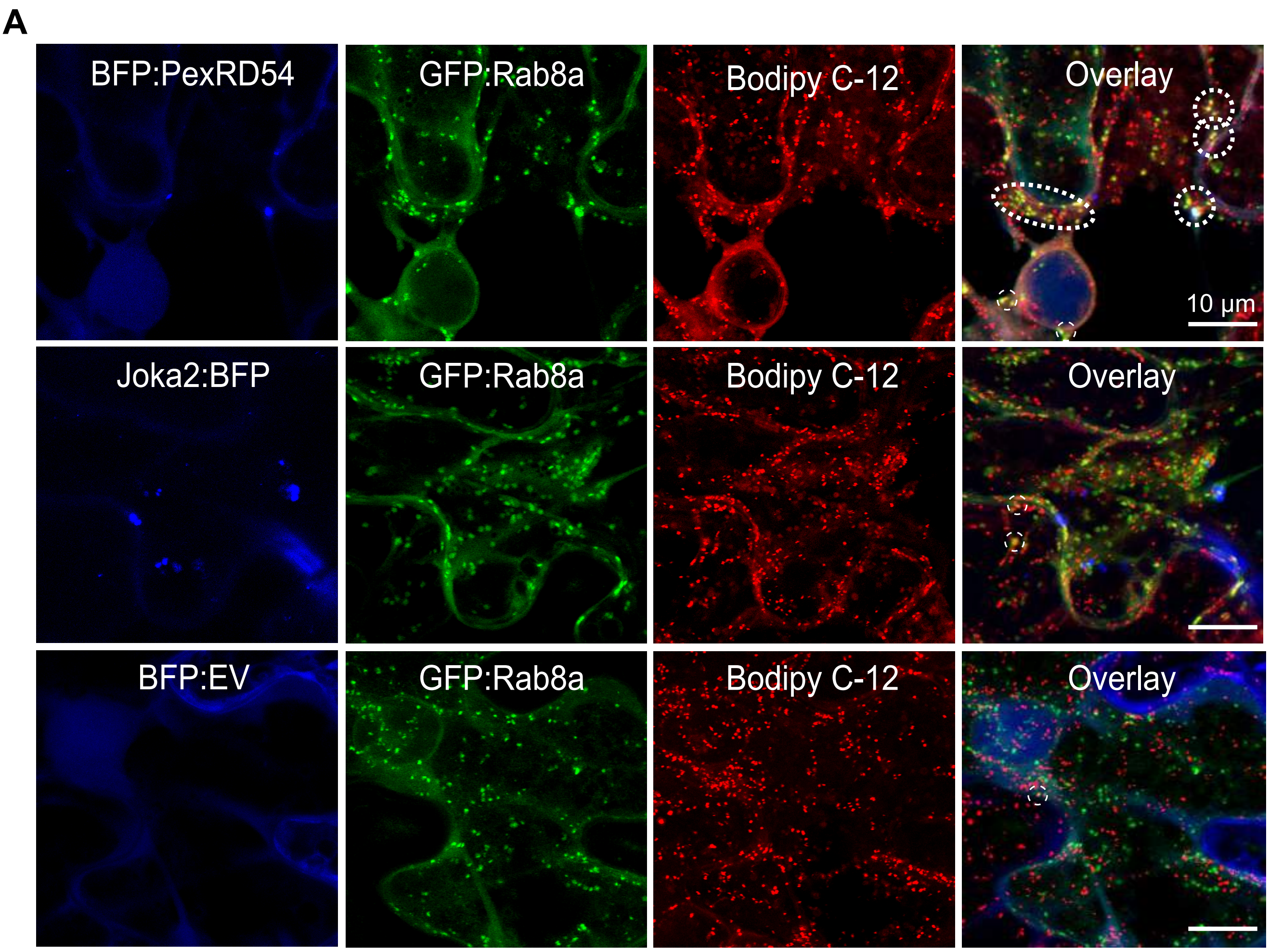


**Figure S25. Stimulation of autophagy by PexRD54, but not Joka2, increases frequency of Rab8a and Bodipy C-12 positive puncta.** Maximum projection confocal micrographs of *N. benthamiana* leaf epidermal cells treated with Bodipy C-12 dye to mark lipid droplets while transiently expressing GFP Rab8 with either BFP PexRD54 (top), BFP Joka2 (middle), or BFP empty vector (bottom).

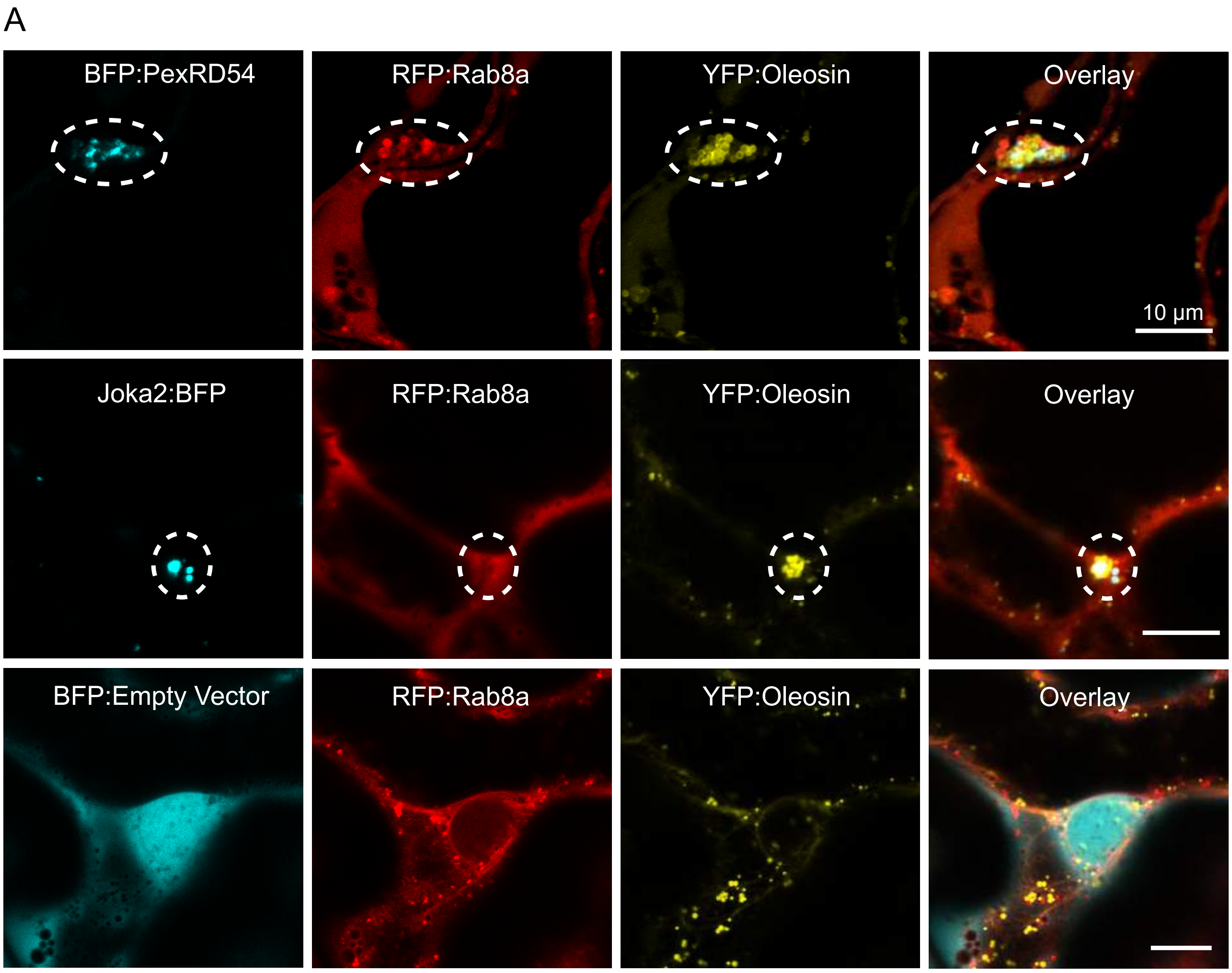


**Figure S26. PexRD54 but not Joka2 recruits Rab8a to Oleosin vesicle-like clusters.** Maximum projection confocal micrographs of *N. benthamiana* leaf epidermal cells transiently expressing RFP:Rab8a, YFP:Oleosin and either BFP:PexRD54, Joka2 or an Empty Vector. Images shown are maximal projections of 15-25 frames with 1 μm steps. Scale bars, 10 μm.

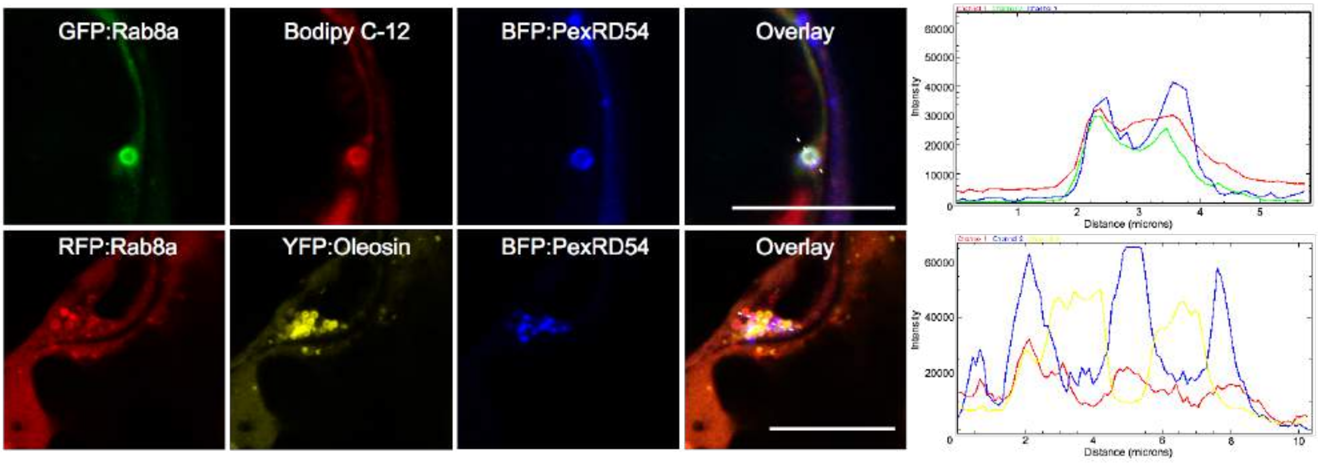


**Figure S27. PexRD54/Rab8a cluster localization with Bodipy C-12 and Oleosin.** Confocal micrographs (xy) of *N. benthamiana* leaf epidermal cells transiently expressing BFP:PexRD54 with either GFP:Rab8a and stained with Bodipy C-12 or RFP:Rab8a and YFP:Oleosin. Transects in overlay panel correspond to plot of relative fluorescence over the labelled distance. Scale bars, 10 µm.

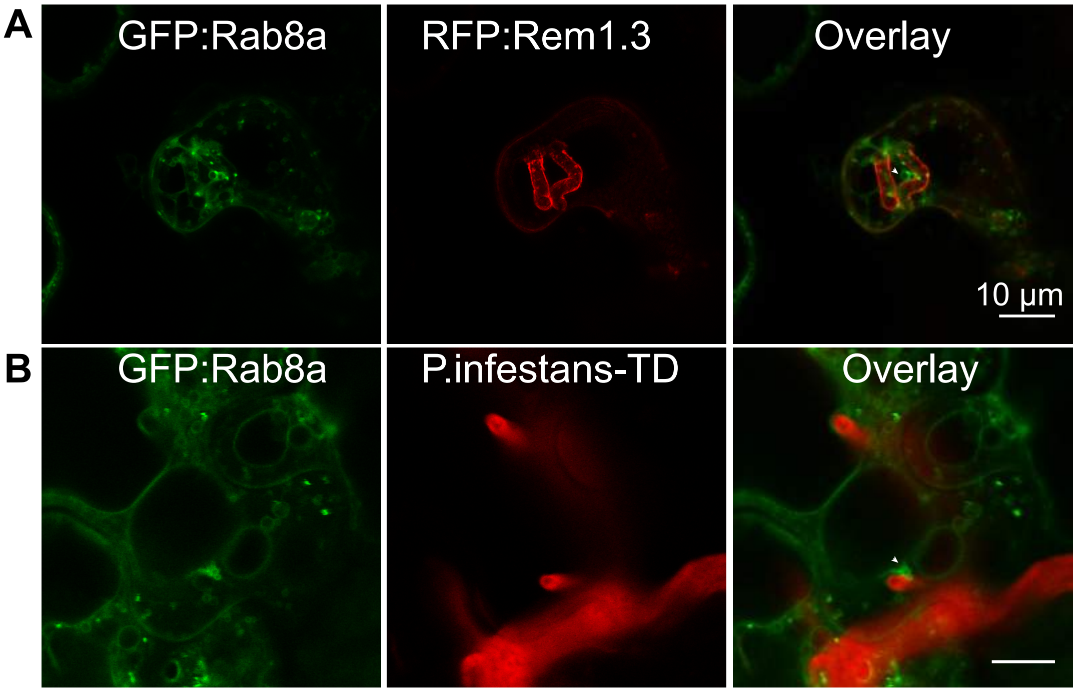


**Figure S28. Rab8a localizes to various vesicle-like structures around haustoria during infection with *P. infestans*.** Maximum projection confocal micrographs of *N. benthamiana* leaf epidermal cells infected with *P. infestans* (3 dpi) and transiently expressing GFP:Rab8a. (A) Tissue infected with WT *P. infestans* and expressing RFP:Rem1.3 to label haustoria. (B) Tissue infected with TD Tomato *P. infestans*. (A, B) White arrowheads highlight *P. infestans* haustoria. Scale bars 10 µm.

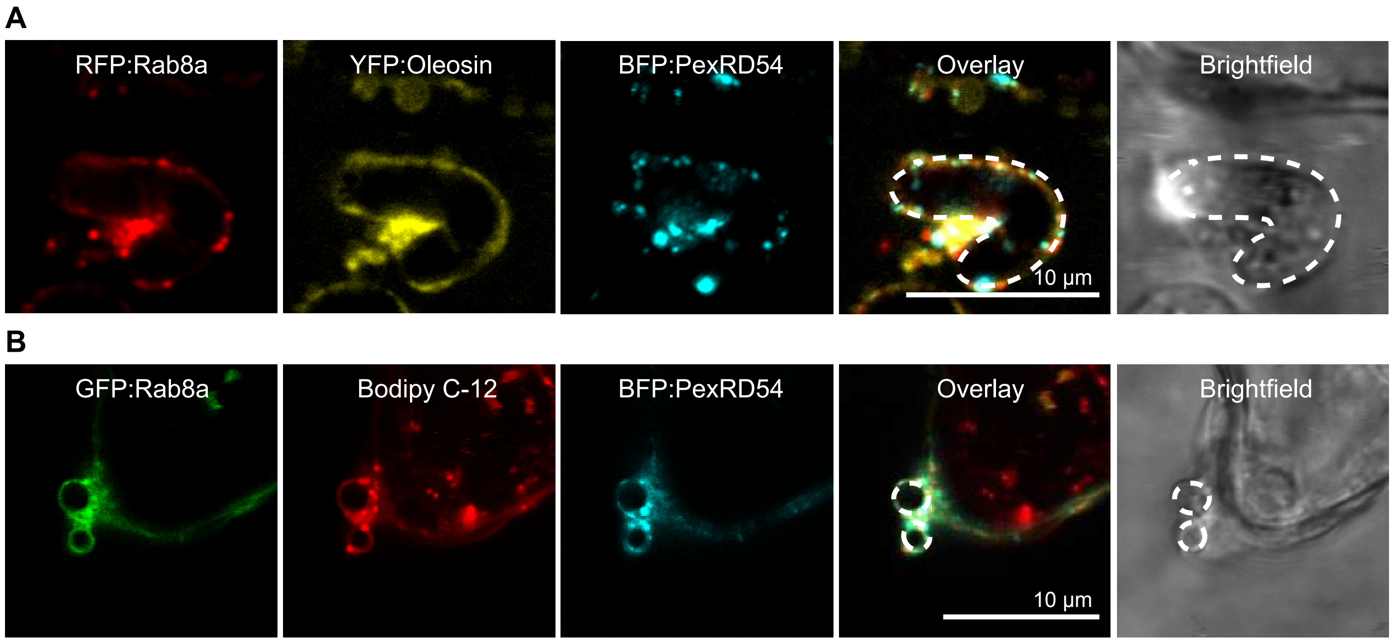


**Figure S29. Rab8a colocalizes with PexRD54, Oleosin and Bodipy C-12 in puncta around *P. infestans* haustoria during infection.** Maximum projection confocal micrographs of *N. benthamiana* leaf epidermal cells infected with *P. infestans* (3 dpi) and transiently expressing (A) RFP:Rab8a, BFP:PexRD54, and YFP:Oleosin or (B) GFP:Rab8a, BFP:PexRD54 and stained with Bodipy C-12. Images shown are maximal projections of 15-25 frames with 1 µm steps. Scale bars, 10 µm. Dotted lines highlight *P. infestans* haustoria.

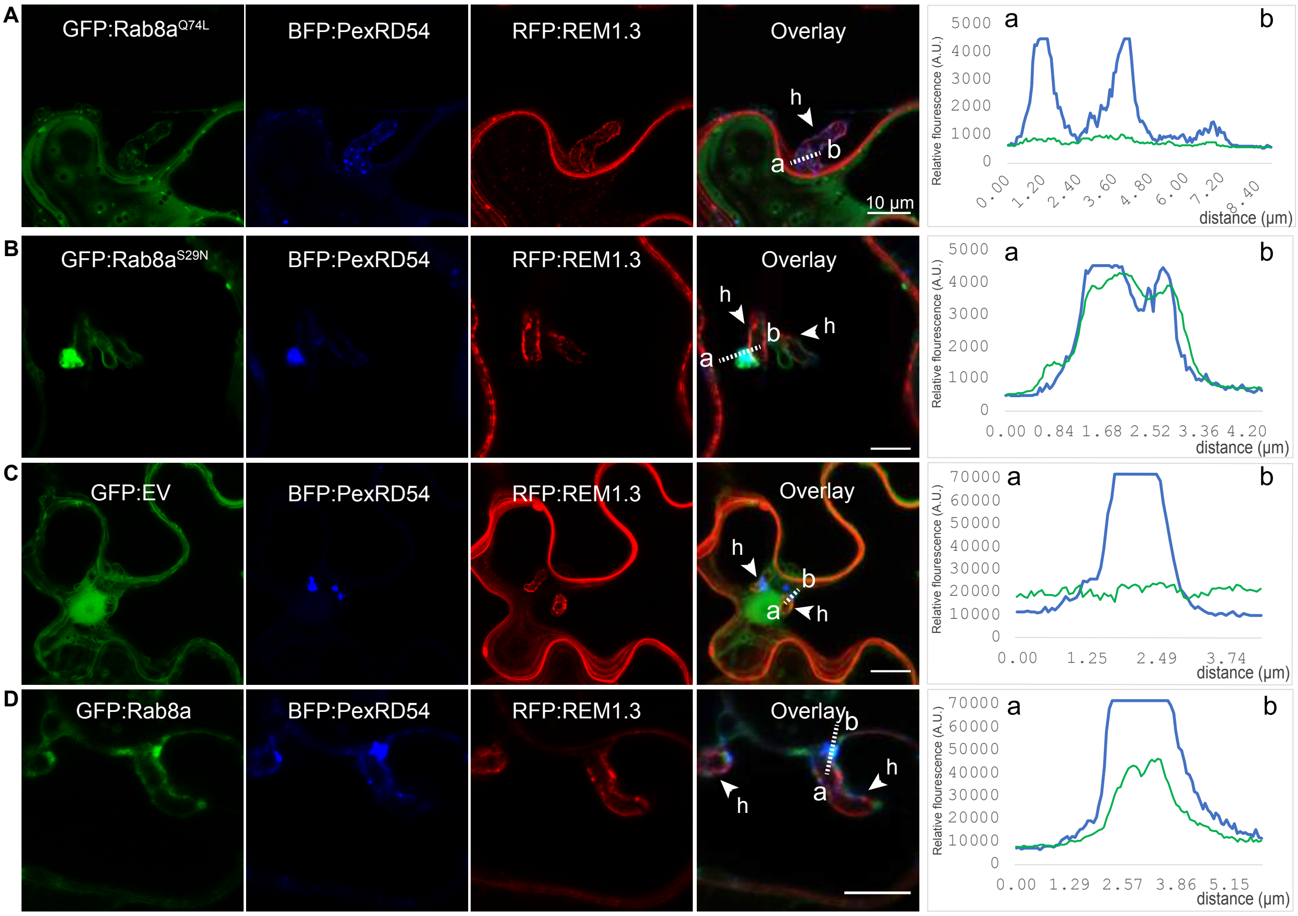


**Figure S30. Localisation of Rab8a and its mutants in perihaustorial autophagosomes during infection by *P. infestans*.** Maximum projection confocal micrographs of *N. benthamiana* leaf epidermal cells infected with *P. infestans* (3 dpi) and transiently expressing either GFP Rab8a GTP (A), GFP Rab8a GDP (B), GFP (C), or GFP Rab8a (D) with BFP PexRD54 and RPF Remorin 1.3. White arrows show haustoria. Transects in overlay panel correspond to plot of relative fluorescence over the labelled distance.

**Video S1. Rab8a localises to small mobile vesicles and large ring-shaped structures.** GFP:Rab8a is co-expressed with the EHM marker RFP:REM1.3 via agroinfiltration in *N. benthamiana* leaf epidermal cells. Confocal laser scanning microscopy was used to monitor Rab8a labelled vesicles three days post infiltration. The movie represents time-lapse of 76 frames acquired during 3 min 48s (Frame interval: 3 s).

**Video S2. Mobile PexRD54/Rab8a positive puncta co-migrate with ATG9 vesicles.** RFP:Rab8a, BFP:PexRD54 and GFP:ATG9 are co-expressed 3 via agroinfiltration in *N. benthamiana* leaf epidermal cells. Confocal laser scanning microscopy was used to monitor PexRD54/Rab8a labelled vesicles and ATG9 compartments three days post infiltration. The movie represents time-lapse of 28 frames acquired during 4 min 12s (Frame interval: 9 s).

**Video S3. Mobile PexRD54/Rab8a positive puncta co-migrate with Bopidy-C12 labelled lipid droplets.** GFP:Rab8a, BFP:PexRD54 are co-expressed 3 via agroinfiltration in *N. benthamiana* leaf epidermal cells stained with the lipid droplet dye Bodipy-C12. Confocal laser scanning microscopy was used to monitor PexRD54/Rab8a labelled vesicles and lipid droplets three days post infiltration. The movie represents time-lapse of 56 frames acquired during 56s (Frame interval: 1 s).

**Video S4. Rab8a localises to small mobile vesicles at haustoria during *P. infestans* infection.** GFP:Rab8a, RFP:ATG8CL and BFP:PexRD54 are co-expressed via agroinfiltration in *N. benthamiana* leaf epidermal cells infected with *P. infestans*. Confocal laser scanning microscopy was used to monitor Rab8a labelled vesicles three days post infection. The movie represents time-lapse of 45 frames acquired during 3 min (Frame interval: 4 s).

**Video S5. Rab8a localises to large vacuole-like structures at haustoria during *P. infestans* infection.** GFP:Rab8a is co-expressed with the EHM marker RFP:REM1.3 via agroinfiltration in *N. benthamiana* leaf epidermal cells infected with *P. infestans*. Confocal laser scanning microscopy was used to monitor Rab8a labelled vesicles three days post infection. The movie represents time-lapse of 26 frames acquired during 26 min (Frame interval: 30 s).
